# Early onset memory deficit of WMI rats compared to their nearly isogenic WLIs is reversed by enriched environment in females

**DOI:** 10.1101/2025.02.20.639157

**Authors:** Michelle T. Ji, Katherine J. Przybyl, Aspen M. Harter, Mariya Nemesh, Sophia Jenz, Anna Yamazaki, Chris Kim, Megan K. Mulligan, Hao Chen, Eva E. Redei

## Abstract

We tested the hypothesis that environmental enrichment (EE) can attenuate early-onset cognitive decline in a stress-hyperresponsive rat strain. The novel genetic model, the Wistar Kyoto More Immobile (WMI) inbred rat strain demonstrates increased stress reactivity and enhanced depression-like behavior compared to its nearly isogenic control, the Wistar Kyoto Less Immobile strain (WLI). Middle-aged (12 months) WMI females exhibited diminished fear, and spatial memory in the contextual fear conditioning and Morris Water Maze paradigms, respectively, compared to young animals (6 months) of both strains and to middle-aged WLIs. Middle-aged WMI males showed a lesser age-induced deficit. EE from six to 12 months of age reversed these memory deficits in middle-aged WMI females and attenuated them in WMI males. Plasma levels of estradiol followed the same pattern as memory in WMI females following EE. RNA sequencing from female hippocampi revealed significant strain, age, and enrichment-induced differentially expressed genes. Among these, solute carrier family 35, member A4 (*Slc35a4*) and potassium inwardly rectifying channel, subfamily J, member 2 (*Kcnj2*) were confirmed to show hippocampal expression changes parallel to that of memory in the WMI. These genes have critical roles in the integrated stress response, cellular metabolism, and the effects of stress on neurovascular coupling, respectively. Pathway analyses revealed the involvement of oxidative phosphorylation and mitochondrial dysfunction in the hippocampal processes of aging and EE-induced reversal. These findings underscore the critical involvement of molecular stress responses in early-onset memory decline and suggest potential therapeutic targets for age-related cognitive impairment.

**Highlights:**

- Heightened innate stress response and depression leads to early onset memory decline
- Enriched environment reverses memory decline of middle-aged females
- Enriched environment also reverses declining estrogen levels of middle-aged females
- Mitochondrial dysfunction may underly mid-life cognitive and molecular changes

## Introduction

The pathological processes underlying dementia, including Alzheimer’s Disease (AD), begin long before clinical onset and last approximately 15–20 years. Therefore, accessing modifiable risk factors that can delay the onset of dementia is critically important (Ren, Liang et al. 2022, Hafizi and Rajji 2023). Stress is among these modifiable risk factors. Stress has been associated with dementia (Livingston, Huntley et al. 2020, Islamoska, Hansen et al. 2021, Luo, Beam et al. 2023), implicated in AD progression, and shown to lower the age of onset (Sotiropoulos, Catania et al. 2011, Islamoska, Hansen et al. 2021, Sulkava, Haukka et al. 2022).

Stress-related disorders like depression have been linked to a state of accelerated aging and subsequent pathological cognitive aging (Herbert and Lucassen 2016, Singh-Manoux, Dugravot et al. 2017, Malhi and Mann 2018, Arnaud, Brister et al. 2022, Sinclair, Mohr et al. 2024). Major depressive disorder (MDD) is a risk factor for mild cognitive impairment (MCI) and dementia, with a prevalence rate of 32% in MCI and up to 37% in dementia (Ismail, Elbayoumi et al. 2017). Even moderate depression increases the risk of progression from healthy to MCI, and to dementia (Kaur, Bucholc et al. 2020).

Studying risk and protective factors for cognitive decline in humans is challenging due to the need to account for genetic, lifestyle, physical, and mental health factors (Unterrainer, Petersen et al. 2024). Deficits in hippocampal cellular function with aging have been shown both in humans and rodents (Bettio, Rajendran et al. 2017). As these deficits start before clinical signs of age-induced cognitive decline in humans, preclinical research models can be useful for the identification of factors influencing early cognitive decline. Few experimental models of early cognitive decline exist. Two well-characterized models include middle-aged Fisher 344×Brown Norway hybrids and insulin-resistant rats (Driscoll, Howard et al. 2006, Stranahan, Norman et al. 2008). Most animal studies show age-induced memory loss in much older animals (Moyer and Brown 2006, Rowe, Blalock et al. 2007, Haider, Saleem et al. 2014, Bettio, Rajendran et al. 2017, Olesen, Torres et al. 2020). Risk factors of cognitive aging such as mid-life or later-life stress are also studied in older animals (Kim and Diamond 2002, Sandi and Touyarot 2006, Borcel, Perez-Alvarez et al. 2008, Arias-Cavieres, Adasme et al. 2017). Stress has variable effects on cognitive aging (Park, Jo et al. 2024), which may be due to genetic variation in stress reactivity and neurocognitive processes.

Although genetic factors contribute to the variation in the risk of AD (Gaiteri, Mostafavi et al. 2016), epigenetic mechanisms are key players in dementia-associated disorders (Jones, Goodman et al. 2015, Delgado-Morales and Esteller 2017, Cui and Xu 2018). The accumulation of epigenetic alterations over the lifespan is suggested to be a mechanism that regulates the transcriptome of aged brain cells affecting learning and memory, and its pathological states (Sweatt 2013, Dias, Maddox et al. 2015, Delgado-Morales and Esteller 2017). Studies of human and rodent hippocampal transcriptomes identified specific changes associated with aging-induced cognitive decline (Park, Valomon et al. 2015, Pereira, Gray et al. 2017, Lanke, Moolamalla et al. 2018). The study by Perreira and colleagues (Pereira, Gray et al. 2017) reveals that most transcriptional changes occur between middle-aged and aged rats, rather than between young to middle-aged animals, again emphasizing the importance of investigating changes in the gene regulatory landscape occurring in aged individuals.

Previously, we demonstrated that female rats from a genetic model of enhanced depression-like behavior and stress-hyperreactivity exhibit premature cognitive decline in middle age (Lim et al., 2018). This genetic model was developed from the Wistar Kyoto (WKY) rat strain, showing depression- and anxiety-like behaviors (Pare and Redei 1993, Pare 1994, Solberg, Baum et al. 2004, Nosek, Dennis et al. 2008). The WKY strain presents traits that mirror several symptoms of major depression (Dugovic, Solberg et al. 2000, Solberg, Baum et al. 2004, Baum, Solberg et al. 2006). Chronic treatments with antidepressants (Jeannotte, McCarthy et al. 2009, Tizabi, Bhatti et al. 2012), electroshock administration (analogous to electroconvulsive therapy) (Kyeremanteng, James et al. 2012) and deep-brain stimulation (Kyeremanteng, James et al. 2012) can all reverse the depression-like behaviors of WKYs. We exploited the fact that the WKY strain was not completely inbred, since it was distributed to different suppliers between the 12-17th generation. Bidirectional selective breeding was conducted using immobility behavior in the forced swim test (FST) as a functional selector (Will, Aird et al. 2003). With subsequent filial breeding, two fully inbred strains were generated: the WMI, and the WLI strains (Andrus, Blizinsky et al. 2012, de Jong, Kim et al. 2021). Each strain differs in response to acute and chronic restraint stress (Will, Aird et al. 2003), in the effects of adolescent stress on adult animals (Kim, Gacek et al. 2021), in affective behaviors (Mehta-Raghavan, Wert et al. 2016, Schaack, Mocchi et al. 2021), in drug intake (Lim, Shi et al. 2018, Sharp, Fan et al. 2021), in stress-enhanced fear learning (Lim et al., 2018; Przybyl et al., 2021), and in premature memory decline (Lim, Wert et al. 2018).

This study aims to extend our previous findings on early memory decline in female WMIs relative to WMI males and to determine whether environmental enrichment (EE) can reverse this deficit. EE has been shown to reverse cognitive deficits associated with aging (Harati, Majchrzak et al. 2011, Simpson and Kelly 2011, Harati, Barbelivien et al. 2013, Cortese, Olin et al. 2018), but its effect on genotype-dependent deficits in early memory has not yet been investigated. Additionally, we quantify changes in the hippocampal transcriptome associated with aging and EE in this genetically defined and naturally occurring model of early onset cognitive loss.

## Materials and Methods

### Animals

Animals were maintained at Northwestern University Feinberg School of Medicine by the Center for Comparative Medicine. All procedures were approved by the Northwestern Institutional Animal Care and Use Committee. Animals were housed in temperature- and humidity-controlled cages, in a 12-hour light-dark cycle, where lights turned on at 0600 hours. Food and water were readily available. The animals used in the study were inbred male and female WLI/*Eer* and WMI/*Eer* rats from the 47^th^-51^st^ generations.

At weaning animals were divided into three groups: standard group housing until 6 months of age (6M) or 12 months of age (12M), and animals that were housed under standard condition up to 6M and then placed into an EE until behavioral testing started at 12 months of age (12M+EE). The EE groups were housed in a larger than standard cage (22 x 14.5 x 8 inch), with plastic toys, chimes, tunnels and chewing toys. All cages were cleaned and changed twice a week and clean toys were provided. The number of animals was 8-11 per group, strain, and sex.

Animals were left undisturbed and then tested in contextual fear conditioning (CFC) at 6 months of age (6M) or 12 months of age (12M and 12M+EE). Two weeks after CFC, animals were observed in the Morris Water Maze (MWM) test. Twenty-four hours after the last component of MWM, animals were euthanized by decapitation within 5 seconds of being removed from their home cage.

Brains and trunk blood samples were collected from study animals. Blood samples were collected in EDTA-coated tubes (0.3µL/0.5mL whole blood, 0.5M). The samples were then centrifuged at 4°C and 4000 RPM for 10 min, and the plasma separated for storage at -80°C. Brains were collected in RNAlater™ (Invitrogen, Carlsbad, CA, USA), a solution that stabilizes and protects cellular RNA, and then stored at -80°C.

### Contextual Fear Conditioning

On day 1, rats were placed into a Technical & Scientific Equipment (TSE, Bad Homburg, Germany) automated fear conditioning apparatus containing a foot-shock grid enclosed by a clear acrylic box. Three minutes of habituation was followed by three mild shocks (0.8 mA, 1 s per min) over a three-minute period. Between animals, the chamber was cleaned using 75% ethanol to eliminate behavioral changes caused by odor. Twenty-four hours later (day 2), the rats were placed in the same chamber for three minutes without any shocks and examined for contextual fear memory via freeze duration and total locomotion (distance traveled). These measures were obtained with a computerized infrared beam system (detection rate 10 Hz) for both day 1 and day 2 of the paradigm. Additional measures collected by the system include the number of rears. Rats that did not respond to the initial shock were excluded from the study.

### Morris Water Maze

After a two-week rest period following CFC, the Morris Water Maze (MWM) test was carried out. The hidden platform version of the test was used as an assessment of spatial learning. MWM was conducted with four consecutive days of learning trials and 6 consecutive trials/day. Trials were recorded and animals were tracked using TSE Videomot 2 software (version 5.75), automatically collecting data for distance, time, and speed in all quadrants of the maze, or near the pool wall for thigmotaxis. A water-filled circular tank (170 cm diameter; water depth 25 cm, temperature 21-22° C) was used. Three visual cues (large black symbols) were placed on the walls surrounding the maze. The platform remained in a constant position 1 cm below water level. Rats were placed in the water in a randomly determined quadrant of the tank. Each rat was allowed to swim either until reaching the platform or until 60 s had passed, at which point it was hand-guided to the platform. Twenty-four hours following the last trial on day 4, the platform was removed, and a 60 s probe trial was conducted, for which animals were placed in the same quadrant opposite the missing platform. Rats were also subjected to a visual platform test on day 5 using a flag attached to the platform. All animals had to locate the visible platform to remain in the study.

Unfortunately, the TSE Videomot software stopped working after 2/3 of the animals had passed through the protocol. Therefore, distance and speed measures could not be collected for all animals, and these measures were not analyzed. The latency to reach the platform measure was scored manually to assure uniformity and compared to the computer-collected measure. The scoring was conducted by an observer blind to strain and group. We found a very high correlation between hand scoring and computer scoring, and therefore the hand-scored latency to reach the platform was used for all groups. Floating or immobility was also scored by an unbiased observer.

### Plasma Hormone Assays

Plasma corticosterone (CORT), testosterone (T), and estradiol (E2) levels were measured by commercially available competitive ELISA kits (Corticosterone Competitive ELISA kit, ThermoFisher, USA; Testosterone ELISA kit, Biomatik, Ontario, Canada; 17-Beta Estradiol ELISA kit, Abcam, Cambridge, United Kingdom) according to the manufacturer’s protocol. The sensitivities of the assays were as follows: CORT, 18.6 pg/ml; T, 49.4 pg/ml; E2, 8.68 pg/ml. Plasma samples were diluted to a 1:1000 ratio for CORT, 1:4 for testosterone, and undiluted for estradiol. The ELISAs were performed in duplicates. ELISA plates were read on the FLUOstar Omega Microplate Reader (BMG Labtech, Ortenberg, Germany). Standard curves were generated using linear regression on log-transformed concentration and absorbance data. The resulting equation was then used to calculate the concentration of the hormones in the samples based on absorbance measures.

### Brain Dissection and RNA Extraction

Brains were thawed on ice and dissected. Hippocampi were dissected on a brain matrix and immediately stored in RNAlater (Invitrogen, Carlsbad, CA) at -80° C. Paxinos rat brain atlas coordinates were used for dorsal hippocampus (AP −2.12 to −4.16, ML 0–5.0, DV 5.40–7.60) and ventral hippocampus (AP −4.20 to −6.00, ML 0–5.00, DV 5.40–7.60) (Wilcoxon et al. 2005).

The dorsal and ventral hippocampal tissue was divided into left and right regions and homogenized using TRI Reagent (Sigma-Aldrich, Saint Louis, MO) and a handheld tissue homogenizer (Kinetica Polytronic). Total RNA was isolated from samples using the Direct-zol RNA MiniPrep Plus kit (Zymo Research, Irvine, CA) according to manufacturer’s instructions. RNA quality was determined using the Nanodrop instrument (Thermo Scientific). Accepted RNA qualities ranged from 1.8-2.2 for the 260/280 and 260/230 ratios. Total hippocampal RNA was derived by combining the separately isolated regions. Samples were stored at -80° C.

### RNA sequencing

Total RNA hippocampal samples (2 strains x 3 conditions x 6 biological replicates; all females) were shipped to Novogene for quality control assessment, library preparation and sequencing. Approximately 50 million paired-end 150 bp reads (range: 42-84 million, mean: 53.6 million) were obtained for each sample. Salmon (Patro, Duggal et al. 2017)was used for the alignment of resulting fastq files to the rat transcriptome (Rattus_norvegicus.mRatBN7.2.107.gtf) and quantification of transcript abundance.

### Reverse Transcription and Quantitative Polymerase Chain Reaction

Reverse transcription was done using the Super Script VILO Master Mix (Invitrogen). Hippocampal RNA (1.0 µg) was used according to the manufacturer’s protocol. qPCR was performed with 5 ng cDNA, specific primer pairs and SYBR Green Master Mix (Applied Biosystems, Foster City, CA, USA), using the QuantStudio 6 Flex Real-Time PCR System (Applied Biosystems). Sequences of primers designed for the target genes are listed in **Supplemental Table 1.** Triplicate reactions were performed for each cDNA sample and analyzed using QuantStudioSoftware (Applied Biosystems). Relative quantification, or RQ values, of target gene expression were determined relative to *Gapdh* and a general cDNA calibrator, acquired from a young adult WLI male, using the 2^-ΔΔCt^ method.

### Data Analysis

#### Behavioral, hormone and gene expression (qPCR) measures

A three-way ANOVA was conducted to analyze the contextual fear conditioning test, plasma CORT levels, and RQ in qPCR across strain (WLI, WMI), age (6M, 12M, 12M+EE), and sex (male, female) using GraphPad Prism version 10.3.1 (GraphPad Software, La Jolla, CA, USA). These analyses were also applied to the probe trial and immobility in the MWM. Plasma T and E2 levels were analyzed by two-way ANOVA in males and females, respectively. All data are presented as mean ± standard error of the mean (SEM). False Discovery Rate (FDR) *post-hoc* analyses were conducted following significant ANOVAs using the two-stage step-up method of Benjamini, Krieger and Yekutieli (Benjamini and Hochberg 2000).

#### Morris Water Maze

Latency data from the Morris Water Maze are positively skewed (i.e., most animals find the platform quickly, while a few take longer) and exhibit greater variance as the mean latency increases. To address these properties of our data, and to evaluate the multiplicative effects of covariates in a more meaningful way, we leveraged the Gamma Generalized Linear Model (GLM) with a log link function to examine the effects of sex, group, strain, and day or trial on day 4 on the latency to reach the platform time. For the primary GLM examining average time across days, the model was fit with 411 residual degrees of freedom (df) and a sample size (N) of 426. The GLM across trials on day 4 was fit with 625 df and N of 641. In this model, exponentiated estimates (Exp_Estimate) represent the multiplicative change in mean latency per unit change in the predictor and are reported with confidence intervals (Lower_CI, Upper_CI) in the results. The primary model included all two- and three-way interactions between sex, group, and strain, as well as the main effect of day, to assess the significance of primary effects and interactions. The same methodology was applied to the data for Day 4 to investigate trial-specific effects. All analyses were performed in R (Team 2023) using the glm function for model fitting. False Discovery Rate (FDR) *post-hoc* analyses were performed as described above for day and trial and are indicated in **Figure 2**. Additionally, the probe trial and floating/immobility data were analyzed by three-way ANOVA, followed by FDR *post-hoc* analyses.

#### RNAseq analyses

Although all RNA samples passed quality testing before library preparation, group differences in RNA Integrity Numbers (RIN) appeared to affect RNA-seq outcomes. Specifically, WMI 12M+EE samples had a higher proportion of lower RIN samples. Therefore, RNA-seq analyses were performed with and without the 12M+EE results. Two subsets of RNAseq data were analyzed that included: (1) the full data set, and (2) a data set excluding the 12M+EE samples from both strains, which exhibited lower RIN values. For subset 1, the raw count data was normalized using the *estimateSizeFactors* function in DESeq2 to correct for differences in library size and contrasts (t-tests) between 6M, 12M and 12M+EE were performed for each strain. A significance threshold of *p* < 0.001 was applied to identify differentially expressed transcripts for each contrast. For subset 2, normalization and differential expression was performed in DESeq2 and contrasts between 6M and 12M were performed for each strain (**Supplemental Tables 2, 3**). In addition, normalized count data was also used to perform a two-way ANOVA (Strain: two levels and Age: two levels) to identify differentially expressed transcripts with a main effect of strain, age, or an interaction effect (**Supplemental Tables 2, 3**). The significance threshold was set at a 10% false discovery rate with a minimum fold change of 1.2. Significant differentially expressed genes (DEGs) from subsets 1 and 2 were used for ontology enrichment analyses.

Pathway and ontology enrichment analyses were performed using Qiagen Ingenuity Pathway Analysis. Please note that we entered the results of RNA-seq into the IPA as human genes, rather than rodent, for better coverage, and that is reflected in the capitalized name of genes.

## Results

### Contextual Fear Conditioning

Contextual fear conditioning in young (6M), middle-aged (12M), and middle-aged enriched environment (12M+EE) WLI and WMI rats confirmed and extended our previous findings (Lim, Wert et al. 2018). While WLIs showed no age-related differences in the freezing response on day 1 of CFC, 12M WMIs exhibited significantly lower freezing compared to both their 6M counterparts and 12M WMIs that underwent six months of environmental enrichment (**Figure 1**; age, F[2,110]=13.41, p<0.001; age x strain, F[2,110]=14.35, p<0.001). There was also a significant sex and strain difference in the effect of aging on fear conditioning on day 1. WMI females at 12M showed a greater decrease (approx. 70%) in freezing behavior following foot-shock relative to 6M females, while WMI males at 12M had a smaller (approx. 35%) decrease compared to 6M males (**Figure 1**; sex x strain, F[1,110]=5.82, p<0.05). No interaction effect was observed in WLIs. No significant main effects of sex and strain were observed during fear conditioning.

**Figure 1.**
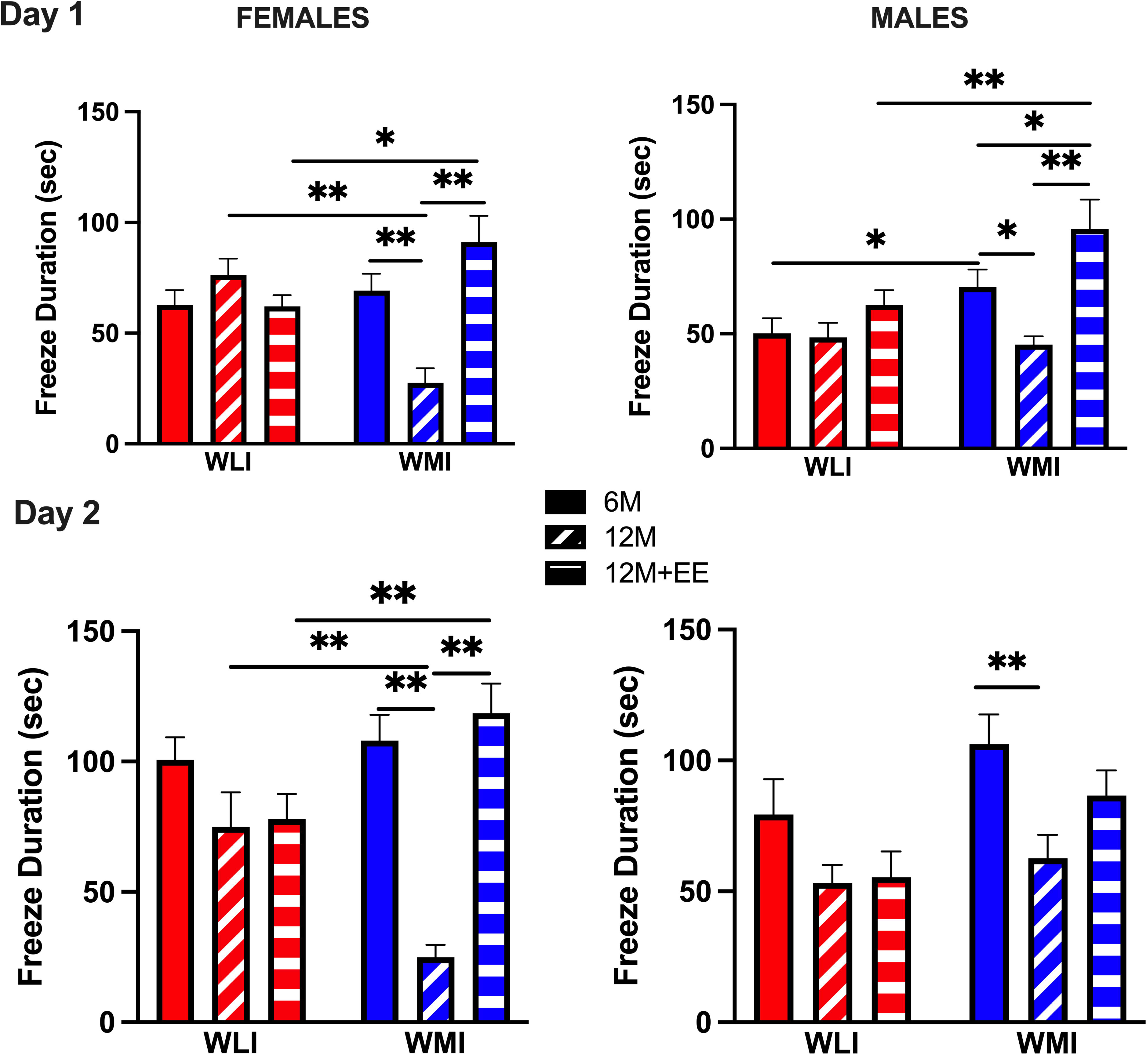
Fear conditioning and fear memory as measured by freeze duration during day 1 and day 2 of CFC. **Day 1.** The overall freeze duration after the three 1 sec foot shock is significantly lower in middle aged (12M) WMIs compared to young (6M) WMIs, and that of middle-aged WMIs after 6 months in EE (12M+EE). **Day 2**. WMI females show decreased fear memory at 12M that is reversed by EE, while 12M+EE WMI males exhibit fear memory that is not different from 12M WMI males. N=8-13/strain/sex/age. Data as mean ± SEM. Statistical differences were determined by three-way ANOVA*. Post-hoc* group comparisons were carried out by two-stage linear set-up procedure of Benjamini, Krieger, and Yekutieli following significant ANOVA *q<0.05; **q<0.01 corrected for multiple comparisons.

Similar to day 1, only 12M WMIs demonstrated significantly attenuated freezing behavior on day 2 compared to 6M WMIs (**Figure 1**; age, F[2,103]=24.21, p<0.001; strain, F[1,103]=5.57, p<0.05; age x strain, F[2,103]=8.97, p<0.01). This age difference between 6M and 12M WMI females in conditioned fear memory on day 2 disappeared after 6 months of enriched environment in 12M+EE WMI females. In contrast, the attenuated fear memory of middle-aged WMI males was not completely reversed by EE (age x sex, F[2,103]=3.64, p<0.05; sex x strain, F[2,103]=7.85, p<0.01; **Figure 1**). Again, the fear memory attenuation was greater in 12M WMI females, close to 80% of fear memory of 6M WMI females, while this was about 50% in 12M WMI males.

Distance traveled during day 1 of CFC did not mirror the freezing behavior, and neither did it parallel distance travelled on day 2 (**Supplemental Figure 1**). Main effect of age and sex indicated generally lower locomotion of 12M+EE animals regardless of strain, that females travelled greater distances than males, and that difference in age effects was also modulated by sex (age, F[2,93]=72.65, p<0.001; sex, F[1,93]=9.64, p<0.01; age x sex, F[2,93]=8.26, p<0.001). The difference in distance traveled by strain was also regulated by age and sex, which was shown by WLI 12M females having decreased locomotion in response to the foot-shocks, while 12M males showed an increase (strain, F[1,93]=5.49, p<0.05; age x strain, F[2,93]=3.20, p<0.05; age x sex x strain, F[2,93]=4.04, p<0.05).

In contrast to day 1, day 2 distance traveled showed the inverse of freezing behavior: an internal confirmation of data validity, specifically in females. WMI 12M females travelled significantly more than either 6M or 12M+EE WMIs, while there was no effect of age on WMI males and WLIs of either sex (age, F[2,92]=18.67, p<0.001; age x strain, F[2,92]=16.79, p<0.001; age x sex, F[2,92]=3.78, p<0.05; sex x strain, F[1,92]=9.25, P<0.01; **Supplemental Figure 1**).

Rearing on day 1 and day 2 of CFC showed patterns not corresponding to either freezing or distance traveled. However, the results confirmed that rearing is not simply a measure of activity (**Supplemental Figure 2**). Female 12M WMIs reared dramatically less than either 6M or 12M WMI females on day 1, with similar behavior observed in WLI females, but not in males of either strain (age, F[2,105]=4.50, p=0.01; age x sex, F[2,105]=7.38, p<0.01). Rearing on day 2 was substantially reduced compared to day 1 and showed an opposite pattern in WLIs and WMIs. Namely, WLI 12+EE females and males reared more than their 6M or 12M counterparts, respectively, while WMI 12+EE females reared less and males showed no differences by age (sex, F[1,106]=9.81, p<0.01; age x strain, F[2,106]=3.29, p<0.05).

### Morris Water Maze

GLM with a log link function was used to examine the effects of sex, group, strain, and day or trial on day 4 on the latency to reach the platform. The model revealed a significant effect of sex (Exp_Estimate = 1.23, CI = [1.10, 1.38], p < 0.001), indicating that females had a 23% longer latency to find the platform compared to males. The model also showed a significant effect of strain (Exp_Estimate = 1.17, CI = [1.03, 1.33], p < 0.05), indicating that WMI animals had 17% longer latencies compared to WLI animals. A significant effect of group was also observed (Exp_Estimate = 1.26, CI = [1.10, 1.45], p < 0.01), with 12M animals exhibiting a 26% longer mean latency to locate the platform than 6M animals. Furthermore, latency significantly decreased across days (Day 2: Exp_Estimate = 0.88, 95% CI = [0.81, 0.95], p < 0.001; Day 3: Exp_Estimate = 0.77, 95% CI = [0.72, 0.83], p < 0.0001; Day 4: Exp_Estimate = 0.72, 95% CI = [0.67, 0.77], p < 0.0001). These estimates indicate that mean latency on Days 2, 3, and 4 were 12%, 23%, and 28% lower than Day 1, respectively. Further post hoc analyses were conducted to explore differential responses dependent on sex, strain, and group (**Figure 2**). Latency to discover the hidden platform is shown for strain and group stratified by sex for all days (**Figure 2A**). The largest differences in latency to find the platform emerged by day 4. 12M WLI females and males had shorter latencies relative to WMIs of both sexes at day 4. Posthoc analysis further revealed significantly longer latencies in both female and male 12M WMIs relative to 6M and 12M+EE WMIs by the fourth day of testing.

**Figure 2.**
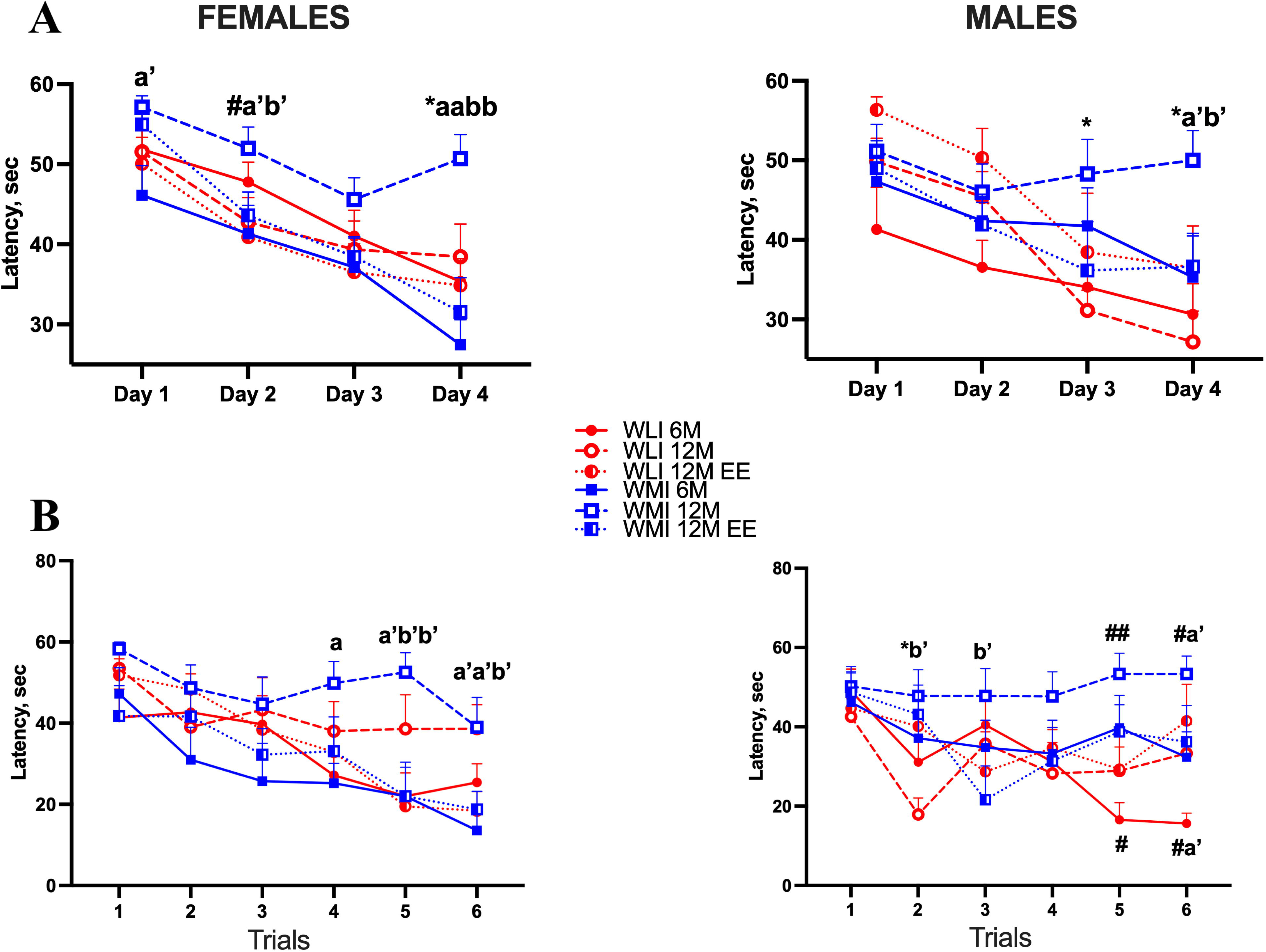
Daily mean latency to reach the platform, and mean latency to reach the platform during each of the six trials on Day 4 in the MWM. **(A)** Six daily trials were carried out over four consecutive days. 12M WMIs took significantly longer to reach the platform especially at Day 4, compared to 12M WLIs and both 6M and 12M+EE WMIs. **(B)** 12M WMI females showed increased mean latency to reach the platform at trials 5 and 6 compared to both 6M and 12M+EE WMI females, while mean latency of 12M WMI males did not differ from 12M+EE WMI males. Values are mean +/-SEM for N=7-10/sex/strain/age. Data were analyzed by Gamma Generalized Linear Model (GLM) with a log link function to examine the effects of sex, group, strain, and day or trial on day 4 on the latency to reach the platform. Post-hoc analyses were conducted by FDR. Significance corrected for multiple comparison are indicated by *, a or b, while p values not corrected for multiple comparison are marked by #, a’or b’. *q<0.05 or #p<0.05, ##p<0.01 shows significance between WLIs and WMIs of the same age group (6M, 12M or 12M+EE) and same day or trial in the MWM; **^a^**q<0.05, **^aa^**q<0.01 or **^a’^**p<0.05, **^a’a’^**p<0.01 represent significance between 12M vs. 6M of the same strain (WLI or WMI) and same day or trial; **^b^**q<0.05, **^bb^**q<0.01 or **^b’^**p<0.05, **^b’b’^**p<0.01 identify significant differences between 12M and 12M+EE of the same WLI or WMI strain and same day or trial. Number of animals as in Figure 1.

A second model was used to examine changes in latency across trials in the final day. Main effects revealed that later trials had lower mean latencies compared to the first trial. Compared to Trial 1, Trial 2 had a 19% lower mean latency (Exp_Estimate = 0.81, 95% CI [0.69, 0.95], p < 0.01), Trial 3 had a 24% lower mean latency (Exp_Estimate = 0.76, 95% CI = [0.65, 0.89], p < 0.001), Trial 4 had a 30% lower latency (Exp_Estimate = 0.70, 95% CI = [0.60, 0.82], p < 0.0001), Trial 5 had a 37% lower latency (Exp_Estimate = 0.63, 95% CI = [0.54, 0.74], p < 0.0001), and Trial 6 had a 39% lower mean latency (Exp_Estimate = 0.61, 95% CI = [0.52, 0.71], p < 0.0001. Further post hoc analyses were conducted in males and females separately to explore differences between group, strain, and trial on day 4 (**Figure 2B**). This analysis demonstrated a more robust effect of group and strain for later trails. Specifically, WMI females at 12M showed increased mean latency to reach the platform at trials 5 and 6 compared to 6M and 12M+EE WMIs. In contrast, 12M WMI males displayed differences from 12M WMI males, specifically, 6M WLI males showed decreased mean latency to reach the platform at trials 5 and 6 compared to 12M and 12M+EE WLIs.

Since the WLI and WMI were selectively bred based on immobility in the FST, we determined that the deficit in recognizing the platform was not related to strain differences in floating (immobility). Floating events during the 6 trials of the last day of MWM were analyzed; every 5 seconds without movement was a floating event. If the animal floated longer than 5 seconds a new event was scored. Data is shown in **Supplemental Figure 3**. Floating was high during the first trial, and it decreased precipitously during the subsequent trials (females: trial, F[5,155]=75.34; p<0.001; males are similar). There were no significant floating/immobility differences between the groups in females (strain, F=0.52, NS; age, F=1.69, NS; interactions, NS). In males, the only significant difference was the decreased floating of WLI at 12M compared to WLI at 6M and WLI at 12M+EE at the first trial (strain, age, NS; trial x age, F[5,90]=10.65, p<0.01).

Twenty-four hours following the last trial on day 4, the platform was removed, and a 60 s probe trial was conducted with animals placed in the same quadrant opposite to the missing platform. The probe trial results showed that 12M WMI females searched for the hidden platform significantly longer before reaching the right quadrant (strain x age, F[2,51]=3.95, p=0.05; sex x age, F[2,51]=7.04, p=0.01; **Supplemental Figure 4**).

### Hormone Measures

Plasma hormone levels were measured in the trunk blood (Figure 3). Plasma CORT levels differed significantly between strains, sexes and by age (strain, F[1,101]=59.48, p<0.001; sex, F[1,101]=11.18, p<0.01; age, F[2,101]=38.88, p<0.001; sex x age, F[2,101]=2.87, p=0.06; strain x age, F[2,101]=19.12, p<0.001; strain x sex, F[1,101]=7.07, p<0.01; strain x sex x age, F[2,101]=2.88, p=0.06; **Figure 3AB**). Analyzing CORT levels of males and females separately revealed that age and strain had a significant effect on plasma CORT levels in females (strain, F[1,46]=8.50, p<0.01; age, F[2,46]=26.87, p<0.001; **Figure 3A**). Middle-aged 12M WMI females had significantly lower levels of CORT than young 6M WMI females. Notably, six months of environmental enrichment increased plasma CORT levels in females of both strains to levels significantly higher than those in their 6M counterparts. This increase was greater in WMI 12M+EE females than in WLI 12M+EE females. In males, only age had a significant effect on CORT levels (F[2,55]=8.94, p<0.001; **Figure 3B**). WMI 6M males showed significantly higher CORT levels compared to WLI 6M males, and EE increased CORT levels significantly higher than those of the young males.

**Figure 3.**
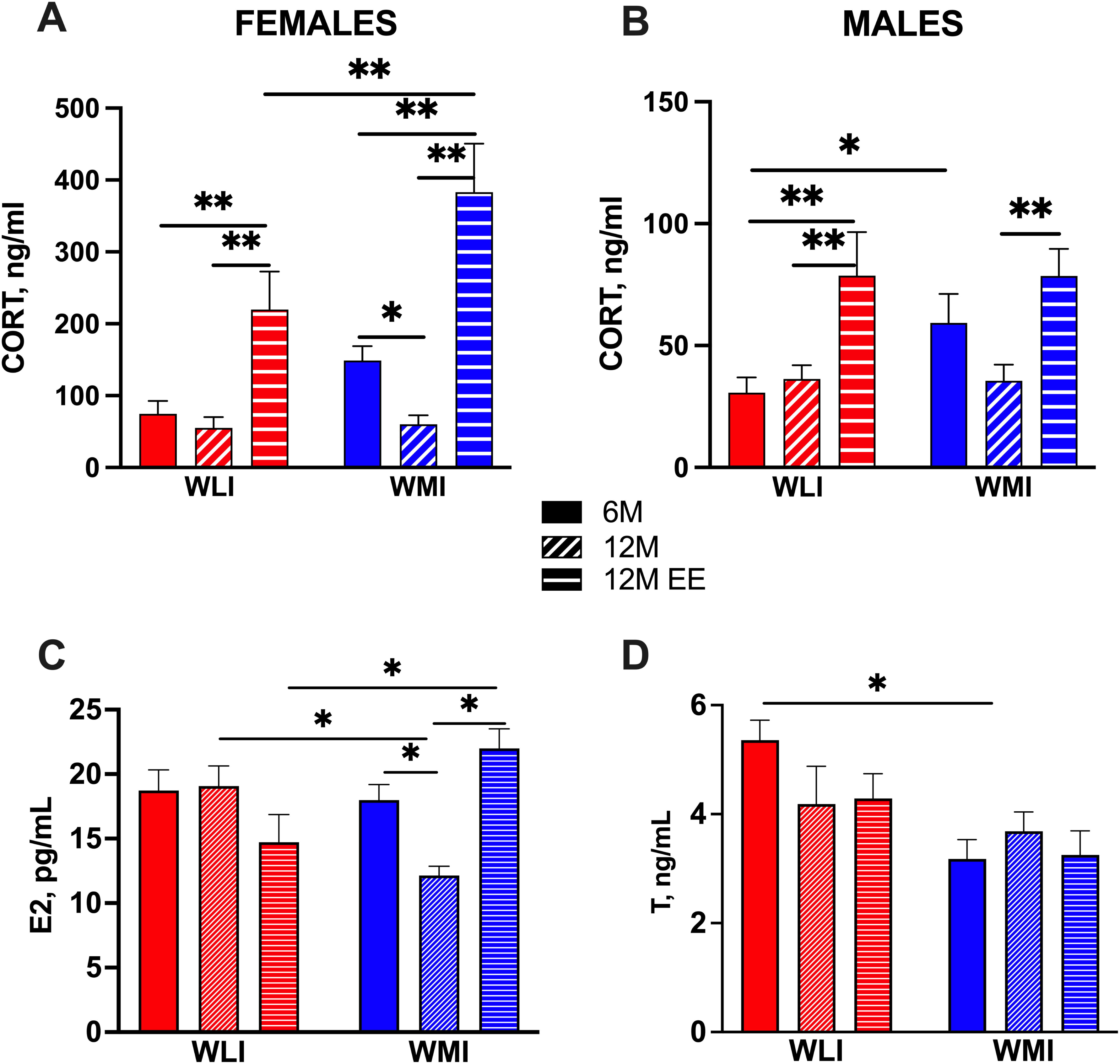
Plasma corticosterone (CORT), estradiol (E2) and testosterone (T) levels from trunk blood. **(A)** Plasma CORT levels were significantly elevated in middle aged females of both strains who spent 6 months in enriched environment (12M+EE). **(B)** In contrast, no significant differences were found in males of either strain by age. **(C)** E2 levels were significantly decreased in 12M WMI females compared to 6M WMIs, but this decrease was reversed by EE in 12M+EE WMI females. **(D)** T levels were significantly lower in 6M WMI males compared to same-age WLI males. Statistics as described in Figure 1. N=8-13/sex/strain/age.

E2 levels did not differ between 6M WLI and WMI females but significantly declined at 12M in WMI females only (strain x age, F[2,52]=7.60, p<0.01; **Figure 3C**). Six months of EE prevented the decrease in E2 in WMI females. In contrast, plasma T levels were already lower in 6M-old WMI males relative to WLI males with no further change due to age or EE (strain, F[2,45]=9.18, p<0.01; **Figure 3D**).

### RNA-seq

Since learning/memory measures and hormone levels changed more dramatically and meaningfully in females, hippocampal RNA sequencing was carried out in female WLIs and WMIs of all three groups.

There were 65 DEGs between 6M WLI and 6M WMI hippocampus (**Supplemental Table S2**). There were substantially fewer DEGs between strains for the 12M comparison, and of the 29 DEGs, only four showed greater expression in the WMI hippocampus compared to that of WLI (**Supplemental Table S3**). We hypothesized that EE would not only reverse memory deficits in 12M WMI females but also restore age-related gene expression changes in the hippocampus. To test this hypothesis, we identified DEGs with an altered expression between 6M and 12M that was reversed by EE in each strain (**Table 1**). In the WLI strain, using a criterion of significance of p<0.0001, we identified 350 DEGs whose expression was decreased by age and increased by EE and 200 DEGs whose expression was increased by age and decreased by EE. For this exploratory analysis we reduced the significance level to p<0.001 for the WMI strain, since the RIN issues (see **Methods**) were primarily present in the WMIs. In the WMI strain, the expression of only 11 and 9 DEGs was decreased by age and increased by EE or increased by age and decreased by EE, respectively.

**Table 1.**
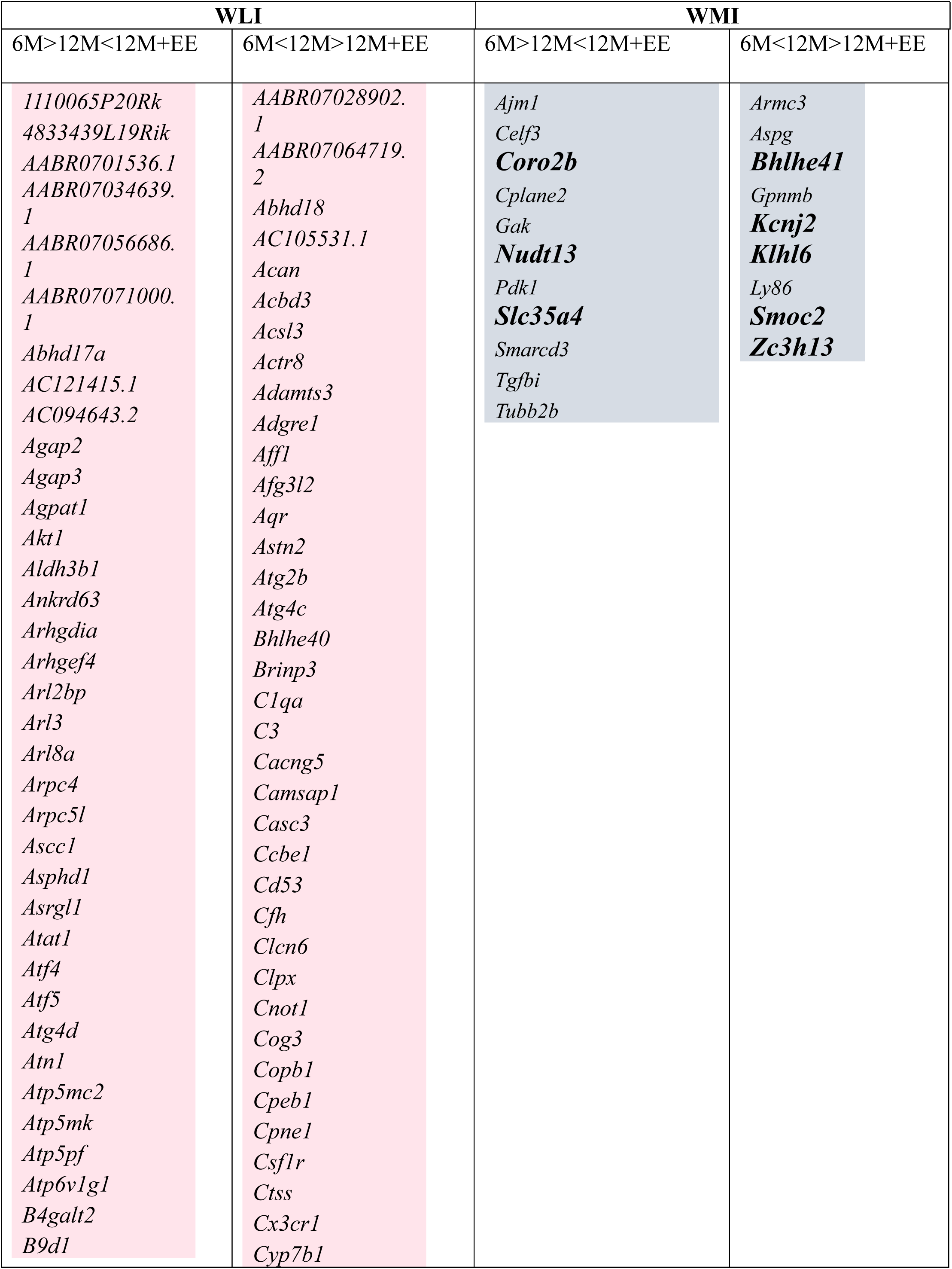
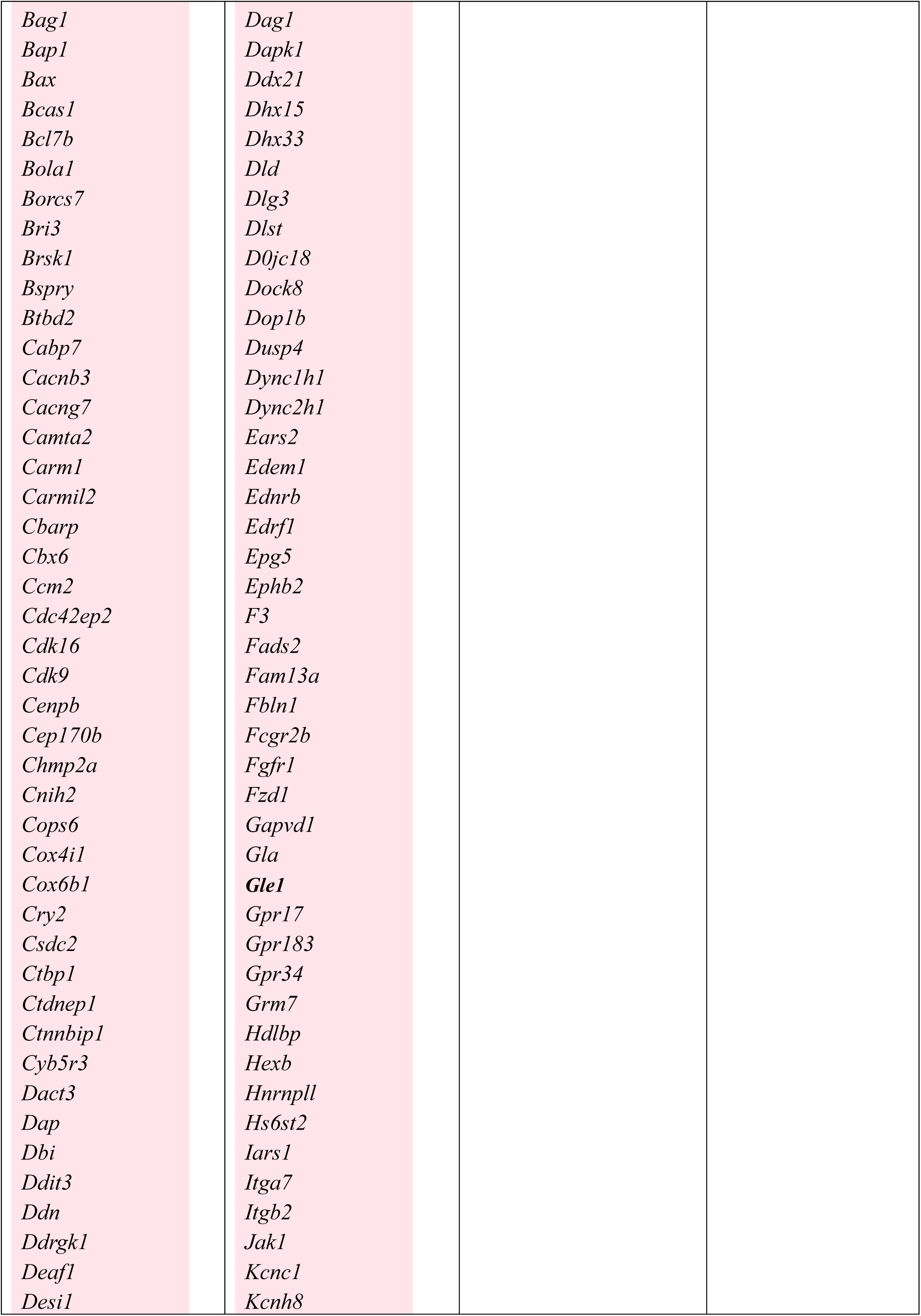
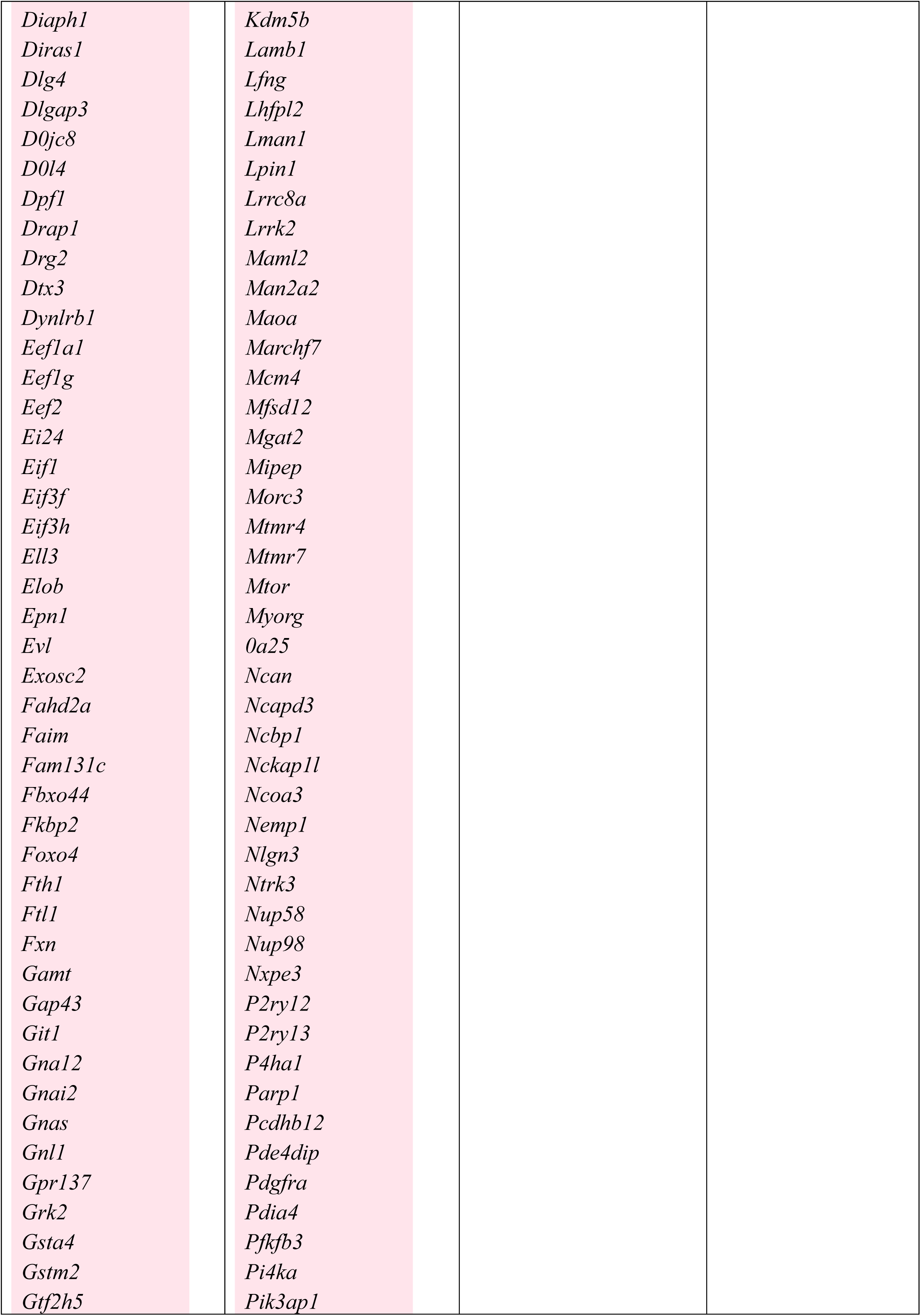
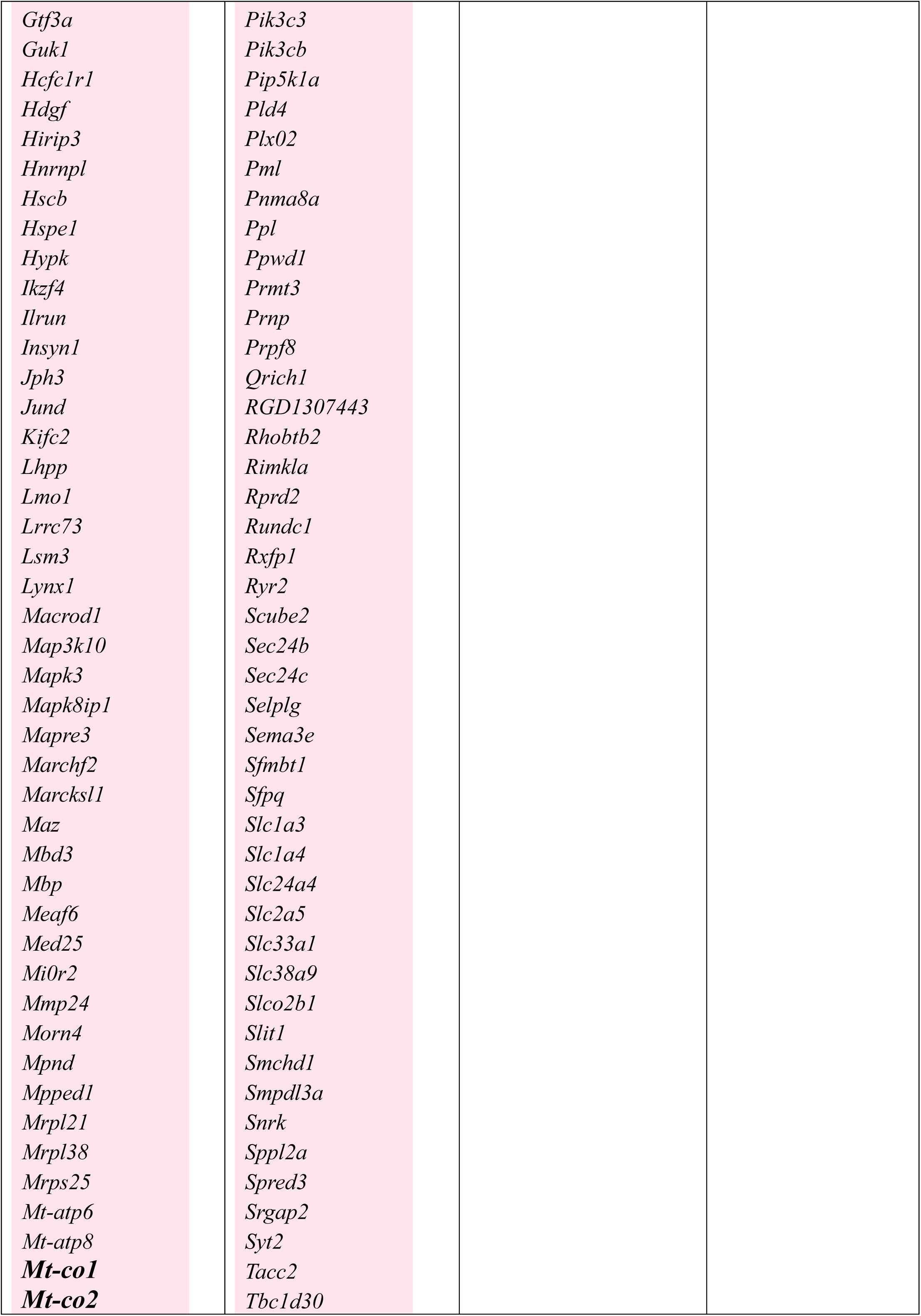
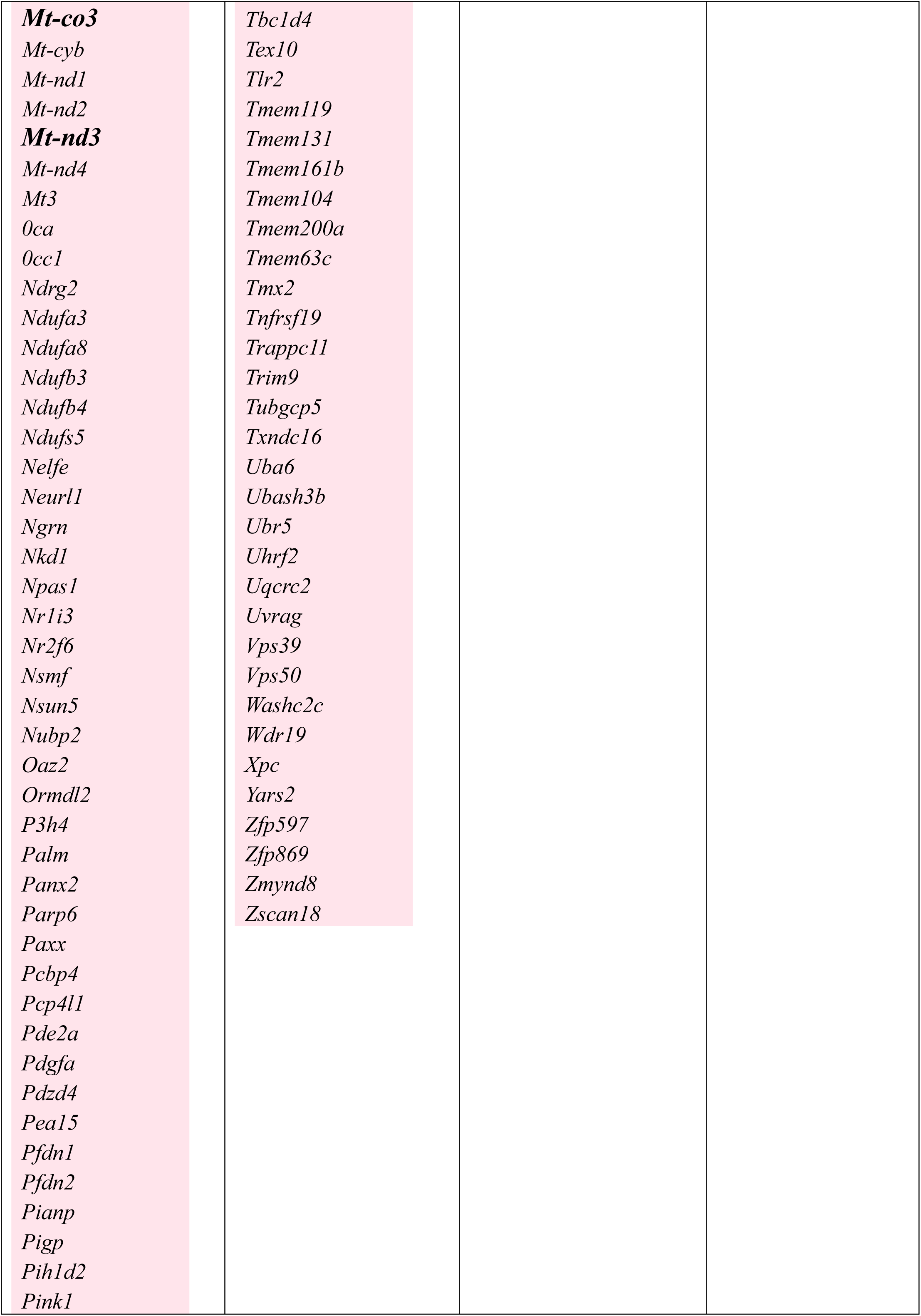
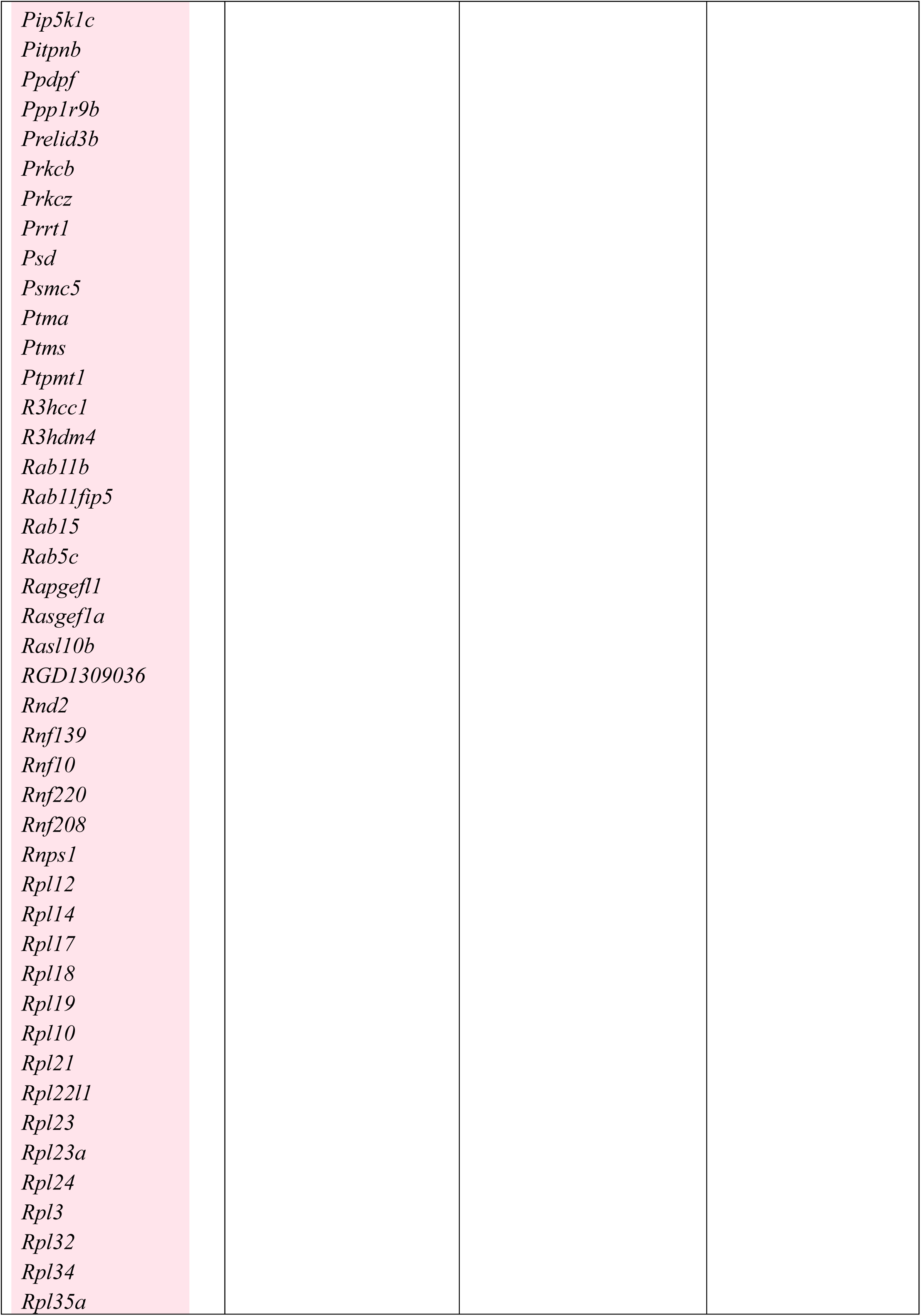
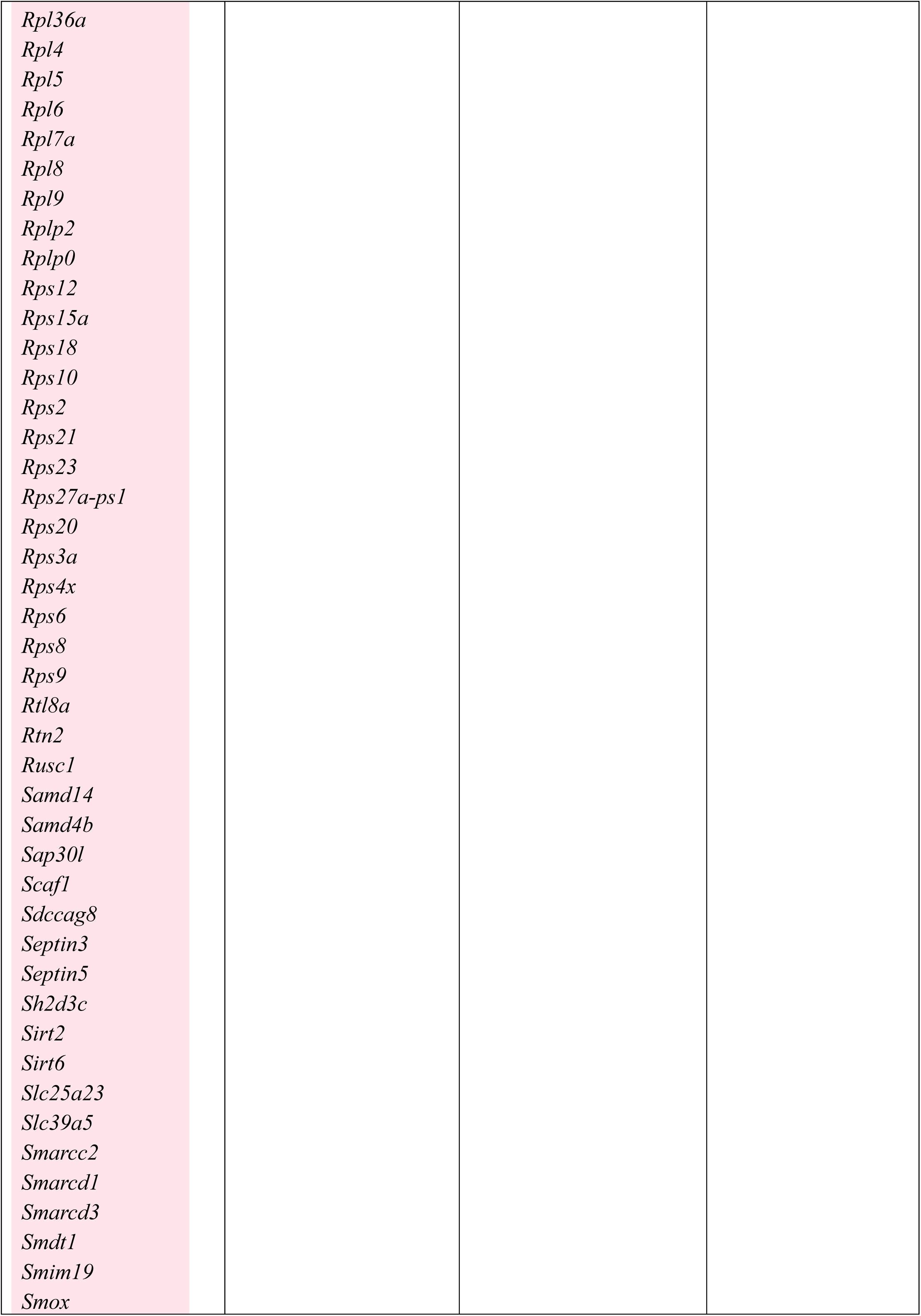
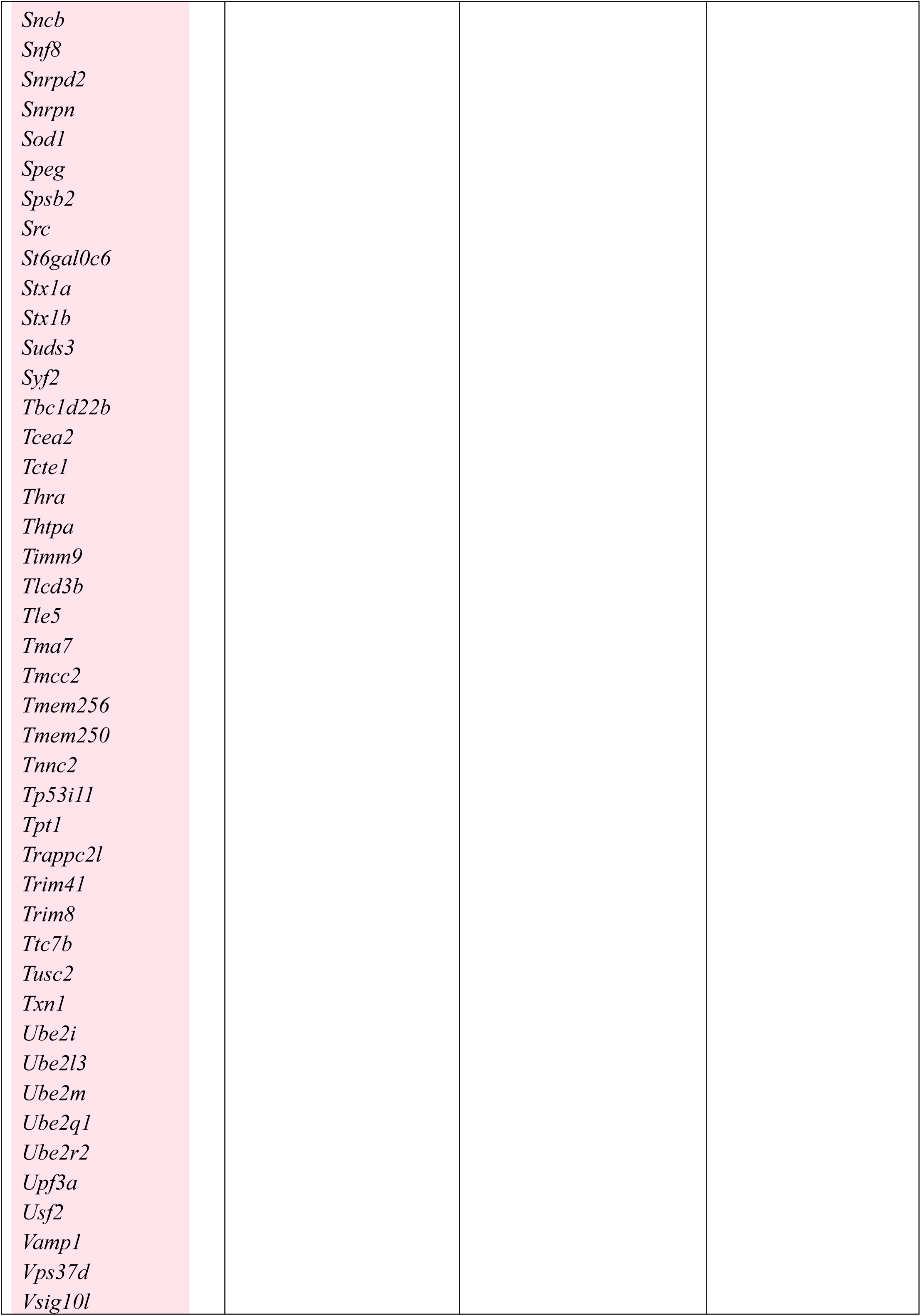
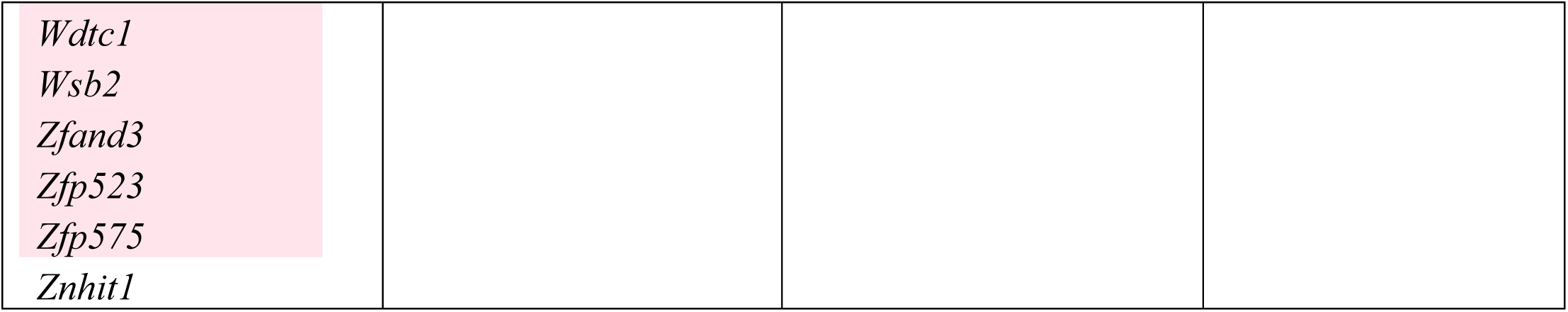
Transcripts with significant (p<0.001) differences between 6M, 12M and 12M+EE.

To verify the WMI-specific DEGs and demonstrate that EE can indeed reverse gene expression changes caused by aging in this strain, hippocampal transcript levels of eight genes (**Table 1 bolded**) were quantified by qPCR in both strains. Significant DEGs were randomly selected to be investigated by qPCR. Result for Coronin 2b (*Coro2b*) and solute carrier family 35, member A4 (*Slc35a4*) confirmed the decreased expression in WMI 12M compared 6M, and the reversal of this decreased expression by EE (*Coro2b*: strain, F[1,44]=3.88, p=0.05; age, F[2,44]=14,01, p<0.001; *Slc35a4*: strain x age, F[1,44]=4.20, p<0.05; **Figure 4AB**). While *Coro2b* expression was also increased in WLI 12+EE compared to 12M hippocampus, *Slc35a4* expression changes were specific to WMIs. In contrast, expression of nudix hydrolase 13 (*Nudt13*), showed no reversal by EE, and opposite patterns between WLIs and WMIs (strain x age, F[1,39]=8.29, p<0.001). Except for *Zc3h13*, all DEGs showing lower expression in 6M than 12M, and a reversal by EE (e.g., lower expression in 12M+EE than in 12M) were confirmed by qPCR. Specifically, hippocampal transcript levels of Kelch-like family member 6 (*Klhl6*), SPARC-related modular calcium binding 2 (*Smoc2*), potassium inwardly rectifying channel, subfamily J, member 2 (*Kcnj2*) and basic helix-loop-helix family, member e41 (*Bhlhe41*) were higher in 12M compared to 6M WMI females, but EE attenuated these increases in the 12M+EE WMIs (**Figure 4**). While expressions of *Klhl6*, *Smoc2* and *Bhlhe41* were higher in WLI 12M hippocampi, and *Bhlhe41* expression also decreased in WLIs by EE, *Kcnj2* transcript level changes were specific for WMIs (***Klhl6:*** age, F[2,35]=18.38, p<0.001; strain x age, F[2,35]=5.25, p=0.01; ***Smoc2:*** age, F[2,41]=13.21, p<0.001; strain x age, F[2,41]=3.13, p=0.05; ***Bhlhe41***: age, F[2,44]=23.15, p<0.001; strain x age, F[2,41]=4.19, p<0.05; ***Kcnj2:*** age, F[2,41]=4.09, p<0.05; **Figure 4**).

**Figure 4.**
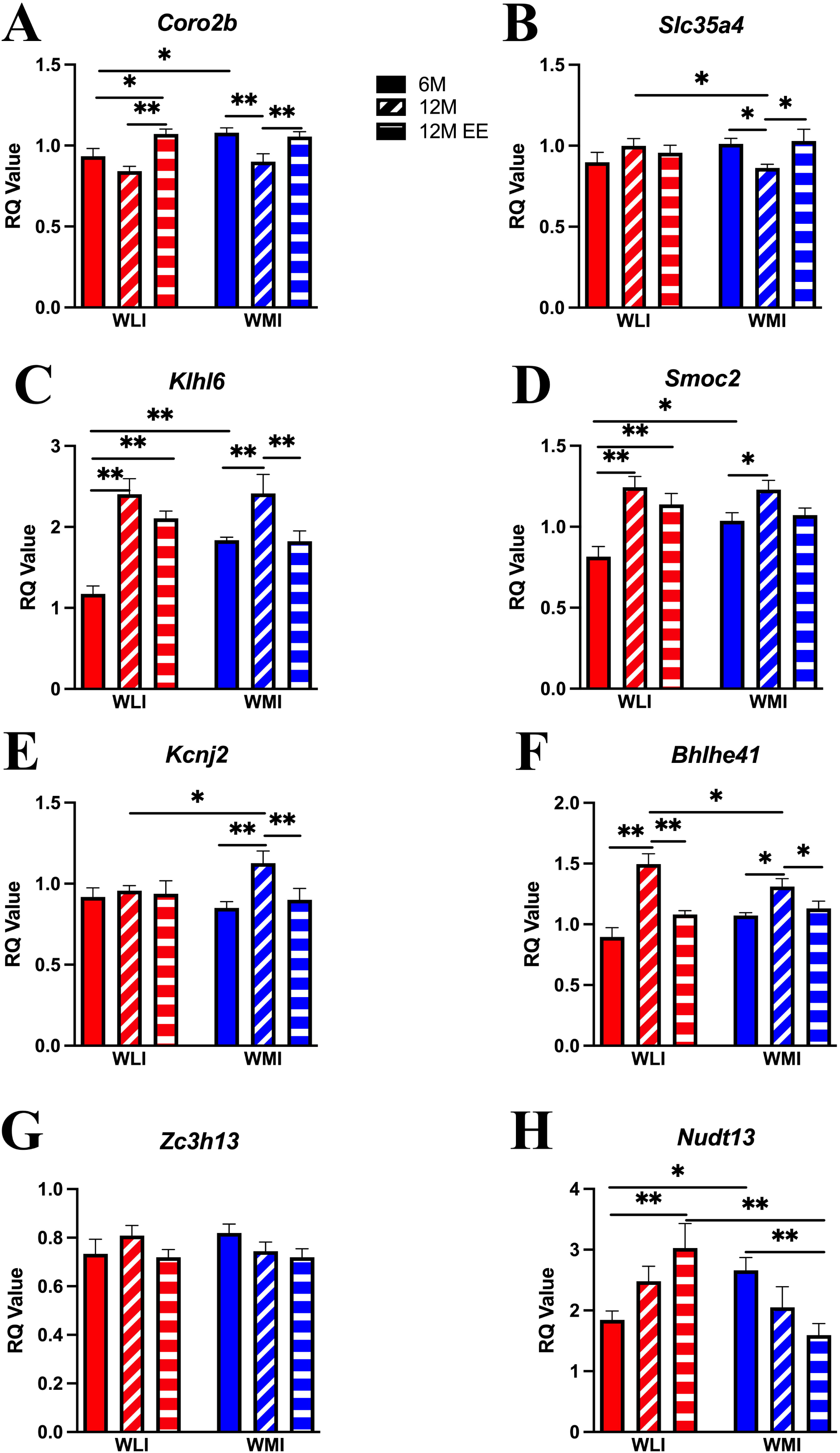
Hippocampal gene expression patterns differ between WLI and WMI females in an age, strain and housing condition-dependent manner. Hippocampal expression of *Coro2b* (A) and *Slc35a4* (B) were significantly lower in WMI 12M females compared to both WMI 6M and 12M+EE WMIs. Hippocampal transcript levels of *Klhl6* (C), *Smoc2* (D) *and Bhlhe41* (F) were higher in both WLI and WMI 12M females compared to 6M levels, but their expression in the 12M+EE hippocampus did not differ from those in 6M females of both strains. In contrast, *Kcnj2* (E) expression only increased in 12M WMI compared to both 6M and 12M+EE WMIs. Transcript levels of *Zc3h13* and *Nudt13* (G, H) did not change by age and strain in contrast to the RNA-seq results. Values are shown as mean **±** SEM. Statistical analyses as described in Figure 1. *q<0.05, **q<0.01, corrected for multiple comparisons. Number of animals as in Figure 1.

The correlations between gene expression quantified by qPCR and RNA-seq data are shown in **Figure 5**. Fold changes for qPCR results were derived from RQs of 6M over RQs of 12M, and 12M +EE over 12M. The correlation between the RNA-seq and qPCR fold changes was significant with r^2^ = 0.756, p<0.001 for 6M vs. 12M, and r^2^ = 0.865, p<0.001 for 12M+EE vs. 12M.

**Figure 5.**
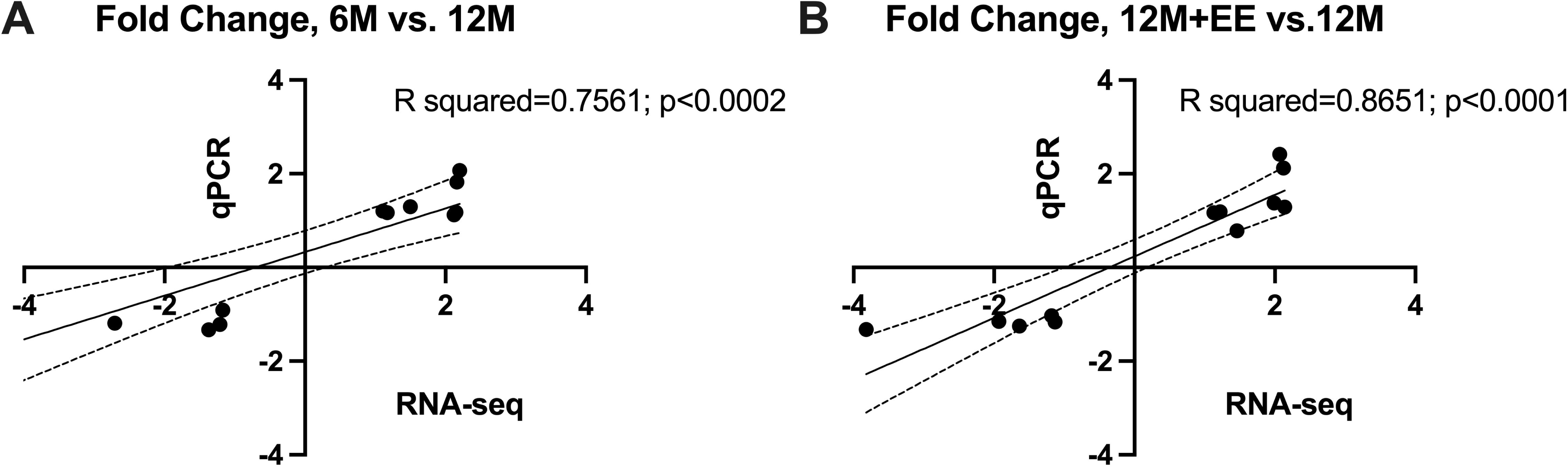
Significant correlation of fold change measures of DEGs between RNA-seq and RT-qPCR. **(A)** Expression of DEGs from RNA-seq, shown in Figure 4, were measured by qPCR in 6M and 12M WLI and WMI female hippocampi. Fold change in qPCR was derived from relative quantification (RQ) of samples. Person correlation between the RNA-seq and qPCR fold changes was highly significant. (**B)** Highly significant Pearson correlation between the qPCR results and the RNA-seq results for 12M vs 12M+EE WLI and WMI female hippocampi.

Significant DEGs were also identified based on age (6M vs 12M) and strain (WLI vs WMI) using a subset of the data that did not include the 12M+EE samples with lower RIN values. Ontology enrichment was used to identify biological processes and pathways represented by these DEGs. Age and strain DEGs were both enriched for the processes of oxidoreduction-driven active transmembrane transporter activity (**Table 2**). Additionally, the overall strain DEGs were enriched for the oxidative phosphorylation term.

**Table 2.** Gene Ontology Enrichment.

| Comparison | Term name | Adjusted p-value | Intersection size |
| --- | --- | --- | --- |
| Age difference (no EE) | protein binding | 8.65e-73 | 2322 |
|  | catalytic activity | 2.03e-22 | 1491 |
|  | molecular adaptor activity | 9.05e-9 | 139 |
|  | channel regulator activity | 1.42E-06 | 56 |
|  | oxidoreduction-driven active transmembrane transporter activity | 1.35E-05 | 24 |
| Strain difference (no EE) | postsynaptic specialization | 4.73e-14 | 20 |
|  | cytoplasm | 8.06e-13 | 92 |
|  | oxidoreduction-driven active transmembrane transporter activity | 5.48e-12 | 10 |
|  | oxidative phosphorylation | 6.96e-11 | 14 |
|  | plasma membrane bounded cell projection | 3.11e-06 | 32 |
| Strain x age interaction (no EE) | protein binding | 5.22e-15 | 389 |
|  | cell adhesion | 1.55e-7 | 81 |
|  | collagen-containing extracellular matrix | 6.07e-7 | 27 |
|  | nucleoplasm | 2.06e-05 | 140 |
|  | cell motility | 7.17e-05 | 88 |
| Overlap between strains by age | phosphate-containing compound metabolic process | 8.19e-05 | 35 |
|  | multicellular organism development | 0.00012 | 49 |
|  | endomembrane system | 0.00013 | 41 |
|  | regulation of protein metabolic process | 0.00027 | 32 |
|  | regulation of amino acid transport | 0.00031 | 6 |

Adjusted p-value is the probability of seeing a number of genes out of the total n genes in the list annotated to a particular GO term, given the proportion of genes in the whole genome that are annotated to that GO term. Intersection size, the number of genes in the input query that are annotated to the corresponding term.

All significant DEGs between WLI and WMI female hippocampus, regardless of age and EE status, were also entered into the Ingenuity Pathway Analysis (IPA) to find the most significant networks (**Table 3**). The top three IPA interactive networks for overall strain differences were characterized as cell morphology, cell signaling, cell-to-cell signaling, developmental disorder, hereditary disorder, metabolic disease, cardiovascular system development and gene expression (**Table 3**). The overall strain differences from data subset 1 that includes the 12M+EE group resulted in the IPA network 2 (score: 39) of “developmental disorder, hereditary disorder, metabolic disease”, as shown in **Figure 6**. The hubs in this network are mitochondrial complex 1, mitochondrial electron transport chain, and cytochrome oxidase processes. The overall age and strain interactive network 1 was categorized into amino acid metabolism, DNA replication, recombination and repair (**Table 3**). Networks 2 and 4 showed the same characterizations of cell signaling, posttranslational modification and protein synthesis, while network 3 was classified by cancer, cell death, survival, organismal injury and abnormalities. The IPA canonical pathway analysis of strain x age interactive DEGs revealed the involvement in oxidative phosphorylation (64 molecules), Eif2 signaling (90 molecules) and mitochondrial dysfunction (107 molecules) (**Table 3**).

**Figure 6.**
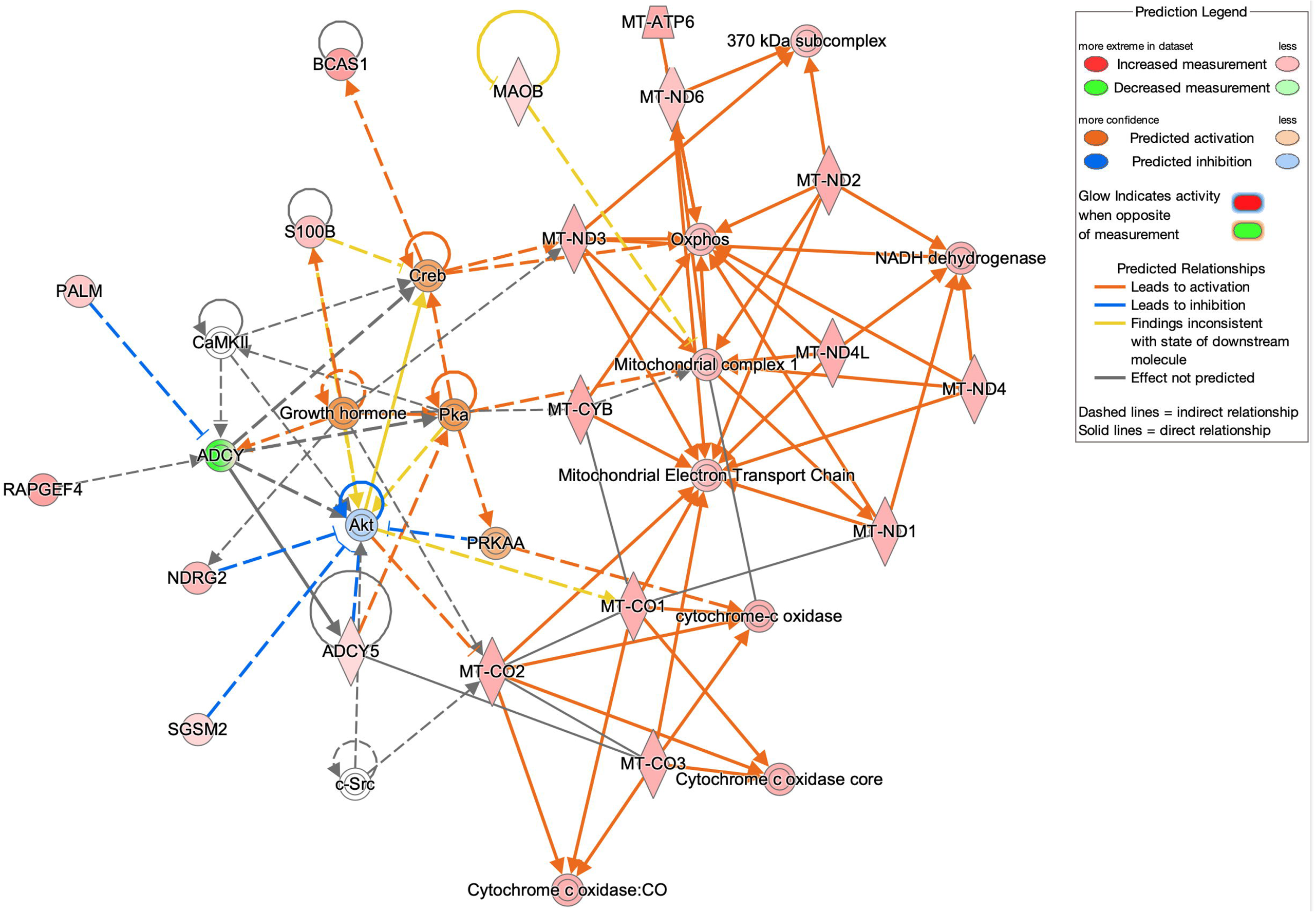
IPA generated network of DEGs from WMI and WLI strain comparison. The IPA network of “developmental disorder, hereditary disorder, metabolic disease”, network 2 (score: 39), is shown. Colored nodes represent input genes. The predicted upstream and downstream effects of activation or inhibition are shown by color overlays and lines, and color coding of the nodes and connecting lines are indicated in the legend. The solid and dash lines denote direct and indirect interactions between two nodes based on the IPA database, respectively.

**Table 3.** IPA networks and canonical pathways.

| **Top IPA networks** |  |  |  |
| --- | --- | --- | --- |
| **Comparison** | **Diseases and Functions** | **Score** | **Focus Molecules** |
| **WLI vs. WMI** | Cell morphology, cell signaling, cell to cell signaling | 42 | 20 |
|  | Developmental disorder, Hereditary Disorder, Metabolic disease | 39 | 19 |
|  | Cardiovascular system development, gene expression | 37 | 18 |
| **Age x Strain** | Amino acid metabolism, DNA replication, Recombination and Repair | 44 | 36 |
|  | Cell signaling, Posttranslational modification, Protein Synthesis | 41 | 34 |
|  | Cancer, Cell death, Survival, Organismal Injury and Abnormalities | 41 | 34 |
|  | Cell signaling, Posttranslational modification, Protein Synthesis | 41 | 34 |
| **Top canonical pathways** |  |  |  |
| **Comparison** | **Canonical Pathways** | **p-value** | **Z score** |
| **Age x Strain** | Oxidative phosphorylation | 1.185E-15 | -7.937 |
|  | Eif2 signaling | 2.7434E-14 | -6.102 |
|  | Mitochondrial dysfunction | 2.0751E-12 | -6.929 |

The oxidative phosphorylation network derived from overall strain differences without EE, the first and third canonical pathways representing DEGs identified from the strain x age (including EE) contrast, and the network hubs in **Figure 6** confirmed the significance of oxidative phosphorylation and mitochondrial dysfunction in molecular aging in WMI and WLI. Based on this, expressions of mitochondrially encoded cytochrome c oxidase 1-3 (*Mt-co1*, *Mt-co2*, *Mt-co3*) and mitochondrially encoded NADH dehydrogenase 3 (*Mt-nd3*) were determined by quantitative RT-PCR in WLI and WMI female hippocampi at six and 12 mos. **Figure 7** shows that aging generally decreased the expression of these genes in both strains, and EE for six months reversed the effect.

**Figure 7.**
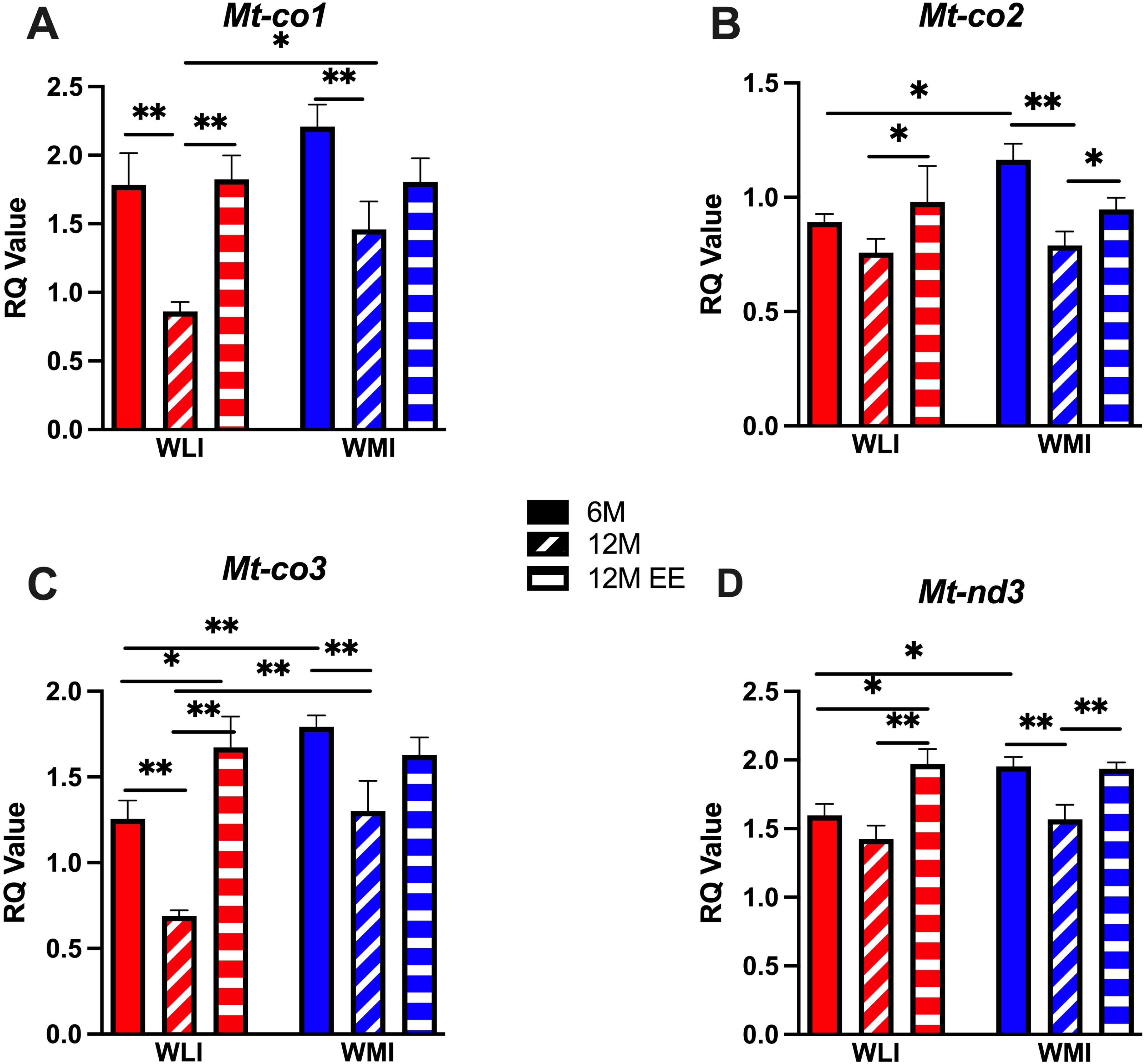
Strain, age and housing conditions affect hippocampal mitochondrial gene expression. **(A)** Decreased expression of *Mt-co1* in 12M female hippocampus in both strains compared to 6M, that is completely or partially reversed by EE in WLIs and WMIs, respectively. **(B) (C) (D)** Enhanced expression of *Mt-co2*, *Mt-co3* and *Mt-nd3* in 6 months old (6M) WMIs compared to WLIs, but EE normalizes these strain differences. *q<0.05, **q<0.01 corrected for multiple comparison.

Expression of these mitochondrial genes was generally higher in WMIs than WLIs, but this difference was eliminated by the increased expression after EE (*Mt-co1*; strain, F[1,42]=5.80, p<0.05; strain x age, F[2,42]=13.57, p<0.001; *Mt-co2*; age, F[2,38]=6.34, p<0.01; *Mt-co3*; strain, F[1,44]=12.35, p<0.01; age, F[2,44]=15.72, p<0.001; strain x age, F[2,44]=3.91, p<0.05; *Mt-nd3*; strain, F[1,45]=4.1, p<0.05; age, F[2,45]=13.16, p<0.001). *Mt-co2* and *Mt-nd3* expression showed greater specificity for WMIs, as the decreased expression at 12M was only reversed by EE in WMIs **(Figure 7)**.

## Discussion

Our study confirms that the genetically stress-hyperresponsive WMI rat, which exhibits enhanced depression-like behavior, experiences early-onset hippocampus-dependent memory loss. Long-term environmental enrichment eliminated this cognitive decline in middle-aged WMI females and attenuated it in WMI males. Analysis of molecular changes associated with the reversal of cognitive decline by EE identified parallel changes in estradiol levels in WMI females. Moreover, RNA sequencing combined with qPCR validation identified genes—*Slc35a4* and *Kcnj2*—whose expression also paralleled changes in memory of WMI females. Furthermore, pathway analysis of DEGs revealed oxidative phosphorylation and mitochondrial dysfunction as key candidate processes underlying hippocampal aging and its reversal by environmental enrichment.

### Memory deficit in middle age

Cognitive changes can occur as early as middle age, 3–7 years before mild cognitive impairment is diagnosed (Karr, Graham et al. 2018). Identifying modifiable risk factors for dementia can facilitate early interventions and reduce disease burden. Literature suggests many modifiable risk factors for dementia (Livingston, Huntley et al. 2020), some of which can be studied in animal models of naturally occurring cognitive decline. Among these are depression and stress. It has been proposed that early treatment of depressive symptoms may impact the course of disease in AD and affect the risk of developing dementia (Dafsari and Jessen 2020). In a large prospective cohort study consisting of over 300,000 participants aged 50-70 years and followed for 10 years depression was linked to a 51% increased risk of dementia (Yang, Deng et al. 2023), reinforcing its role as a significant risk factor. Even subsyndromal depression is associated with cognitive decline (Zhang, Wei et al. 2020).

Stress is another major risk factor, and psychological stress in adulthood is shown to be associated with an increased risk of dementia (Franks, Bransby et al. 2021). Although stress is a very elusive construct, it is thought to be associated with 20-30% higher risk of dementia (Barak 2022). Generally, stress-related neuropsychiatric disorders such as post-traumatic stress disorder (PTSD), general anxiety disorder, and panic disorder exacerbate age-related cognitive decline (Yehuda, Engel et al. 2005, Beaudreau and O’Hara 2008, Greenberg, Tanev et al. 2014) and double the risk of developing AD and other dementia in older individuals (Qureshi, Kimbrell et al. 2010, Yaffe, Vittinghoff et al. 2010). Does depression and stress accelerate cognitive decline starting in middle age? Indeed, there is evidence that stress, stress-related disorders, and depression can accelerate age-related cognitive decline (Han, Schnack et al. 2021, Roberts, Liu et al. 2022).

### Strain differences

A key finding of this study is that the WMI strain, characterized by enhanced depression and stress reactivity, exhibits early-onset cognitive decline compared to the WLI control strain. We use the term early onset cognitive aging in the context of the available literature on aging animal models. Rats usually live for 24-36 months, and most aging studies are carried out with rats over 18 months of age (Ianov, De Both et al. 2017, Haberman, Monasterio et al. 2019). For example, aged Sprague-Dawley rats (>24 months old) exhibit deficits in contextual fear conditioning, whereas late middle-aged (16–24 months old) or early middle-aged rats do not (Moyer and Brown, 2006).

However, the process of cognitive decline can start earlier than the above age range as suggested by the increased heterogeneity of aging-induced cognitive decline observed in late middle-aged (16-20 months old) rats (Haberman, Monasterio et al. 2019). Impaired episodic/working memory emerges around middle-age (reviewed in (McQuail, Dunn et al. 2020) and continues to decline with advanced age (Ando and Ohashi 1991, Markowska and Savonenko 2002, Dellu-Hagedorn, Trunet et al. 2004, Sabolek, Bunce et al. 2004, Templer, Wise et al. 2019, Febo, Rani et al. 2020). The WMI strain showed deficits in fear memory and in spatial memory as early as 12 months of age compared to their nearly isogenic control strain, which is strong evidence of differential genetic and epigenetic contributions to early memory deficits between strains.

### Sex differences

Early-onset memory loss was more pronounced in middle-aged WMI females than in WMI males, contradicting findings from some studies. Deficits in recall after contextual fear conditioning emerge only after 8 months of age in *APPswe/PS1ΔE9* female mice, in contrast to male mice where the recall deficits are already seen at 2 months of age (Kommaddi, Verma et al. 2023). In contrast, non-transgenic studies suggest a male advantage in the effect of aging on spatial working and reference memory for rats across strains (Jonasson, 2005). This sex difference may explain why most studies on aging-induced cognitive decline focus on males (Ianov, De Both et al. 2017, Haberman, Monasterio et al. 2019).

In contrast, our naturally occurring and non-transgenic model of age-induced cognitive decline mimics the sex difference that rules the human aging literature; human females are more vulnerable to cognitive aging than males (Brookmeyer, Evans et al. 2011, Levine, Gross et al. 2021, Guo, Zhong et al. 2022). The dramatic early onset memory loss in middle-aged WMI females, compared to WMI males and WLI females, is demonstrated by a greater loss of fear memory and the inversely exaggerated activity in the CFC paradigm. The freezing response to the conditioning foot shock stimuli paralleled the fear memory in middle-aged WMIs, suggesting that there is also an impairment in the learning process that is more pronounced in the WMI females. Similarly, middle-aged WMI females showed impaired spatial learning in the MWM throughout the four days of training, manifesting in greater latency to reach the platform compared to young WMIs.

It is plausible that the decreased plasma estradiol levels could play a causative role in the attenuation of learning and memory in middle-aged WMI females. Estrogen deficiency heightened the learning and memory deficit in the APP/PS1 triple transgenic mice (Luo, Zeng et al. 2022). Estrogen deficiency induces hippocampal apoptosis and cognitive dysfunctions (Djiogue, Djiyou Djeuda et al. 2018, Yagi and Galea 2019), while estrogen treatment, including brain-specific estrogen prodrug, improves learning-memory performances (Esperanca, Stringhetta-Villar et al. 2024, Salinero, Abi-Ghanem et al. 2024). Thus, reduced estrogen levels in middle-aged WMI females, compared to young WMIs, and unchanged E2 levels in middle-aged WLI females parallel both fear and spatial memory measures in females. Testosterone levels in males were consistently lower in WMI males than in WLI males, regardless of age, thus age-induced attenuated memory in WMI males does not parallel changes in peripheral testosterone levels.

### Environmental enrichment on stress, depression and memory

Long-term environmental enrichment reversed the early onset memory loss, specifically in WMI females. Environmental enrichment enhances learning and memory while mitigating age-related cognitive decline in animals (van Praag, Kempermann et al. 2000, Frick, Stearns et al. 2003, Bennett, McRae et al. 2006). The main mechanism by which EE affects behavioral and molecular mechanisms is that it can attenuate the negative effects of stress by either acting on the same neurological pathways concurrently or acting on different pathways in parallel (Macartney, Lagisz et al. 2022). Additionally, EE is beneficial for reducing anxiety and depression and increasing cognitive performance (see (van Praag, Kempermann et al. 2000), and stress can negatively impact the same traits (Sandi 2004, Macartney, Lagisz et al. 2022). It remains unclear whether EE mitigates aging effects by counteracting stress in stress-reactive WMIs. However, plasma CORT levels were increased, as opposed to decreased, in both WLIs and WMI females after EE, and there was no significant effect of EE on memory measures of middle-aged WMI males. Thus, the specificity of EE in reversing the cognitive decline of middle-aged WMI females, could be related to restoring plasma E2 levels in WMI females. How that increase in E2 levels occur in the presence of elevated plasma CORT is not clear. Nevertheless, this increase could affect oxidative stress and mitochondrial function, both of which are known to be involved in age-induced cognitive decline (Yaribeygi, Panahi et al. 2018, Butterfield and Halliwell 2019).

Lower serum levels of E2 are observed in female patients with AD compared to appropriately matched controls (Barron and Pike 2012). Estrogen has been shown to improve cognitive functions and alleviation of depression in humans and rodent models (Hwang, Lee et al. 2020, Luine and Frankfurt 2020). In a recent study, reduced transcript levels of estrogen receptors (ESR) were found in postmortem brain regions of female subjects with AD and major depressive disorder (MDD), compared to those with AD and no MDD (Luo, Pryzbyl et al. 2022). Since hippocampal ESR2 expression correlated with transcript levels of antioxidant enzymes in that study, females with AD could have exaggerated accumulation of oxidative stress because of the reduced estrogen-induced mitochondrial defect, compared to controls (Long, He et al. 2012). Thus, it is feasible that estrogen-regulated processes contribute to the vulnerability to early onset cognitive decline in WMI females.

### Hippocampal transcriptomics

While aging is linked to cognitive decline, individual variability in age-related memory loss exists in both humans and animals (Bettio, Rajendran et al. 2017, Haberman, Monasterio et al. 2019). The hippocampus is of particular interest for aging and cognitive decline as it is known to play an important role in learning and memory consolidation. For these reasons, we quantified hippocampal gene expression in females as a first step towards identifying the underlying mechanisms through which EE can improve memory function in WMI females.

We hypothesized that EE would reverse age-related gene expression changes, which was confirmed by the restoration of hippocampal *Slc35a4* and *Kcnj2* expression. Interestingly, the SLC35A4 protein encoded by this gene is thought to regulate sensitivity to oxidative stress, as a knockout of *SLC35A4* enhanced sensitivity to oxidative stress in a rescuable manner, indicating direct involvement of SLC35A4 in stress resistance (Ajala, Tiaiba et al. 2024). Additionally, *SLC35A4* codes for an inner mitochondrial membrane microprotein crucial for cellular respiration and ATP generation with a vital role in cellular metabolism (Rocha, Pai et al. 2024).

Likewise, *Kcnj2* (KIR2.1) encodes an inwardly rectifying potassium channel expressed in glial cells whose function has been found to be impaired following oxidative stress, leading to neuronal hyperexcitability, a situation that can be reversed by antioxidant treatment (Remigante, Spinelli et al. 2024). Additionally, the elevation of K+ in the perivascular space activates smooth muscle Kir channels to cause vasodilation and increased blood flow in response to the activity of the nearby neurons (Filosa, Bonev et al. 2006). In rats, this activity driven vasodilation is impaired by chronic stress via glucocorticoid receptor-dependent downregulation of Kir 2.1 (Longden and Nelson 2015). In human subjects, acute psychosocial stress elicits changes in hemodynamic response in insular, temporal, and prefrontal cortices. These hemodynamic changes were associated with genetic differences in *KCNJ2* expression (Elbau, Brucklmeier et al. 2018). In agreement with these findings, WMI 12M females showed reduced corticosterone levels (**Figure 3)** and increased *Kcnj2* expression compared to their 6M controls (**Figure 4**). Both of these changes were reversed by EE, therefore, one additional potential mechanism of EE is remediating the impaired hemodynamic responses to stress in the brain.

Both *Slc35a4* and *Kcnj2* seem to respond to oxidative stress and its alleviation. Moreover, the expression of both genes is differentially altered (*i.e.*, decrease in *Slc35a4* and increase in *Kcnj2*) as a function of age in the middle-aged WMI female hippocampus, and these changes in expression were reversed by EE. Taken together, these genes are likely to contribute to the enhanced vulnerability to aging-induced cognitive decline via altering sensitivity to oxidative stress, which may be reversed through EE.

Substantiating this hypothesis were our findings that mitochondria and oxidative stress pathways were enriched in gene lists based on age and strain interactions, which could be the results of age-aggravated excess of oxidative stress in WMI females. We have previously reported differential vulnerability to oxidative stress and mitochondrial dysfunction between strains even at a very early age (Ferreira, Harter et al. 2024); embryonic WMI neurons and astrocytes are more vulnerable to oxidative stress compared to WLI . Although the expression of many mitochondrial genes was altered by aging and reversed by EE in the WLI strain (**Table 1**), the low number of DEGs detected in the WMI strain is likely due to low RIN values, especially in the 12M+EE condition. Moreover, when the expression of mitochondrial genes was quantified by qPCR in independent samples (**Figure 7**), we found that these genes are more highly expressed in WMI relative to WLI females at 6M, expression is reduced at 12M in both strains, and EE restores expression to about the same level in both strains.

### Study Limitations

Although we confirmed that EE reverses age-related changes in hormones, gene expression, and cognitive decline in middle-aged WMI females, the underlying causal mechanisms remain unclear. EE can act through multiple pathways that ultimately lead to changes in stress reactivity, depression, cognitive function, metabolic function, and estrogen levels. Transcriptome analysis was limited by technical issues (e.g., low RINs), which were more prevalent in the WMI experimental group. Despite this limitation, we were able to validate many of the changes in gene expression, including in mitochondrial genes, *Slc35a4* and *Kcnj2,* but we were unable to detect many more exclusive changes in the WMI strain because of technical issues. Thus, our gene expression data alone cannot resolve whether some or all these potential underlying mechanisms are responsible for the cognitive protection afforded by EE treatment in the WMI strain.

In addition, although we detected changes in mitochondrial gene expression associated with strain, aging, and EE, future studies will be required to interpret the contribution of these changes to cognitive function. Specifically, quantification of mitochondrial abundance and function will be required as an increase in mitochondrial gene expression could indicate either an increased number of mitochondria or an increase in oxidative stress/mitochondrial metabolic burden, both of which can have a divergent impact on cognitive function.

### Conclusions

In this paper, we confirmed the early onset decline of hippocampal-dependent memory in the stress and depression-prone WMI rat relative to its nearly isogenic control strain, the WLI. Importantly, we demonstrate that an intervention during adulthood, EE, can reverse cognitive decline in midlife in the WMI strain, with more profound effects in female WMIs relative to males. To better understand the underlying molecular mechanisms that might be driving genetic background-by-age-associated cognitive decline, we profiled hippocampal gene expression in females, and plasma CORT, E2, and T levels in both sexes at different ages and under different environmental conditions.

We identified and validated gene expression changes associated with aging that were reversed by EE, some of which (*i.e.*, *Slc35a4* and *Kcnj2*) showed greater specificity for WMIs. We also noted the strong effect of EE on mitochondrial gene expression, possibly with a more beneficial effect in the WMI strain. Finally, we identified E2, but not CORT or T, as a potential mediator of EE’s beneficial effect on memory in WMI females. Although we were unable to identify the causal molecular mediators underlying the profound therapeutic effect of EE on cognitive decline in our model, our work advances the field in at least two critical ways.

First, we establish the nearly isogenic WMI and WLI strains as a novel model for investigating cellular and molecular interactions among aging, interventions, and genetic vulnerability to cognitive decline. Second, we propose candidate genes and testable hypotheses regarding the molecular mechanisms underlying genotype × age × EE interactions, focusing on E2 levels and hippocampal expression changes in mitochondria and oxidative response genes. Future work leveraging forward genetics approaches, such as genetic mapping to identify underlying variants in age-associated cognitive decline, combined with epigenetic profiling and functional assays to determine vulnerable cell populations and precise alterations in mitochondrial number, function, and gene expression are expected to reveal molecular processes underlying the age-associated decline in memory between strains. A forward genetics approach is especially useful in the context of the WMI/WLI model based on the very limited number of sequence variations between the two strains (de Jong, Kim et al. 2021).

## Author Contributions

*Conceptualization:* Redei EE, Mulligan KM, Chen H, Ji TM; Przybyl KJ. *Investigation/Data curation:* Ji TM, Przybyl JK, Harter MA, Nemesh M, Yamazaki A, Kim C. *Analysis and interpretation of data:* Ji TM, Przybyl JK, Harter MA, Jenz S, Mulligan KM, Chen H, Redei EE. *Writing-original draft:* Ji TM, Redei EE, Mulligan KM, Chen H. *Writing-review and editing:* Ji MT, Przybyk KJ, Harter MA, Nemesh M, Jenz S, Yamazaki A, Kim X, Mulligan KM, Chen H, Redei EE; *Funding acquisition:* Chen H, Mulligan KM, Redei EE, Ji TM

## Acknowledgments

This work was supported by the Davee Foundation to EER, a grant from the Northwestern University Office of Undergraduate Research, Weinberg College of Arts and Sciences to MTJ and NIH grant R01DA048017 to HC, MKM, EER.

## Statement

During the preparation of this work the authors used GPT-4o to proof the final draft for grammatical errors and clarity. After using this tool/service, the authors reviewed and edited the content as needed and take full responsibility for the content of the publication

## Supplemental Figure Captions

**Supplemental Figure 1. Distance traveled in the CFC was not correlated between day 1 and day 2, but distance traveled on day 2 was inversely related to fear memory in WMIs. Day 1.** The overall distance traveled after the three 1 sec foot shock was significantly lower in middle aged (12M) compared to young (6M) females of both strains. This activity was further decreased by EE in females. 12M+EE males of both strains traveled less than their young and middle-aged counterparts. **Day 2.** Contrary to day 1, distance traveled was significantly and solely higher in 12M WMIs compared to both 6M and 12M+EE WMI males and females. 8- 13/strain/sex/age. Data as mean ± SEM. Statistical differences were determined by three-way ANOVA. Post-hoc group comparisons were carried out by two-stage linear set-up procedure of Benjamini, Krieger, and Yekutieli following significant ANOVA *q<0.05; **q<0.01 corrected for multiple comparisons.

**Supplemental Figure 2. Rearing during CFC. Day 1.** Rearing events showed an age, strain and sex-dependent pattern like that of distance traveled. **Day 2.** In general males reared more than females and EE affected rearing to the opposite direction in WLI and WMI females. Statistics as in Supplemental Figure 1.

**Supplemental Figure 3. Floating/immobility during the last day of Morris Water Maze test.** There were no significant differences by age, strain and housing condition in floating across the trials in males and females.

**Supplemental Figure 4. Probe trial latency to target quadrant on day 5 of Morris Water Maze test.** The platform was removed on day 5 and the animals placed into the water maze at the opposite quadrant to the target quadrant. Almost all 12M WMI females did not find the quadrant during the available 60 seconds.

