## Supplementary figures and images for "Early onset memory deficit of WMI rats compared to their nearly isogenic WLIs is reversed by enriched environment in females"

### Supplemental Figure 1

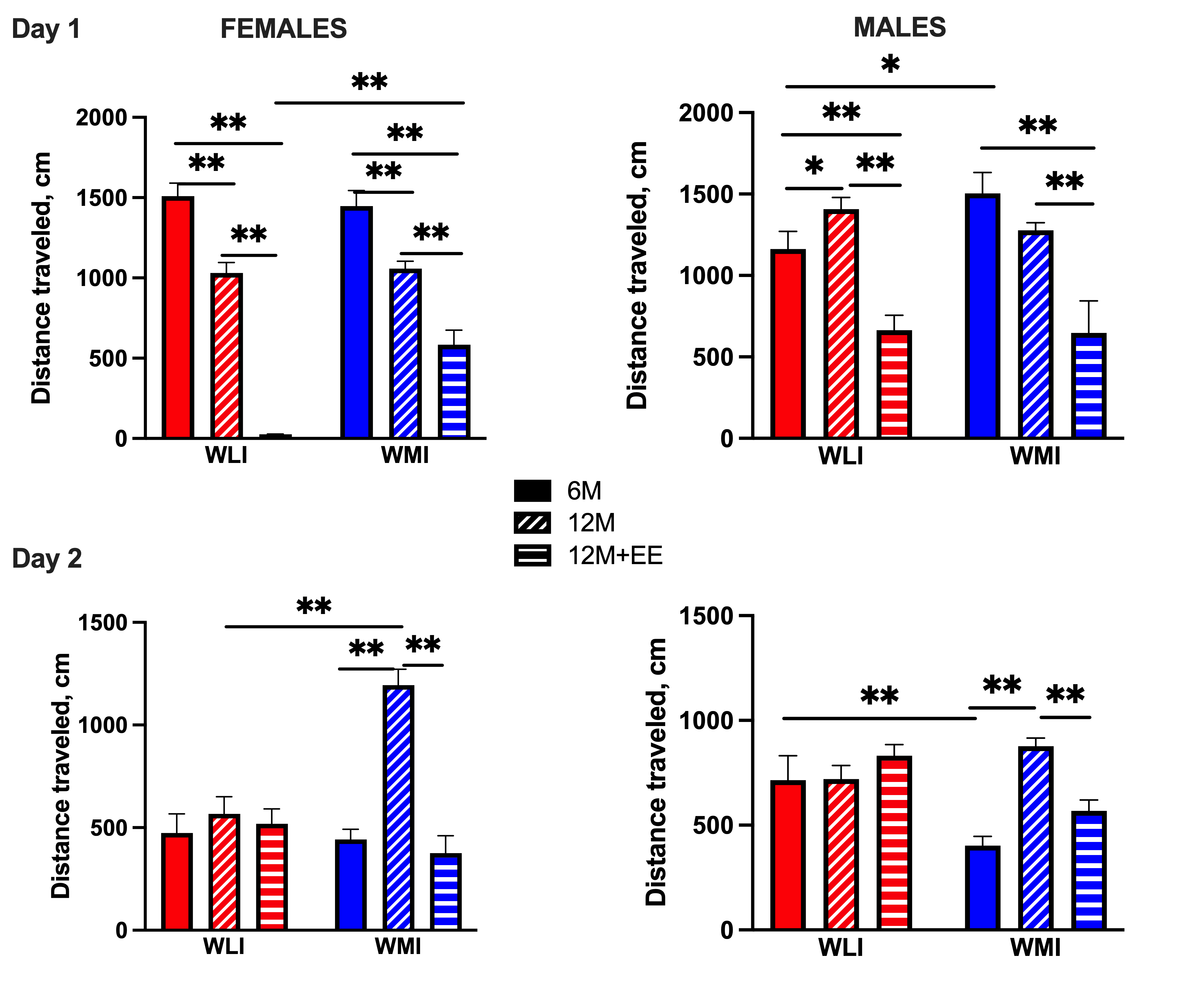

### Supplemental Figure 2

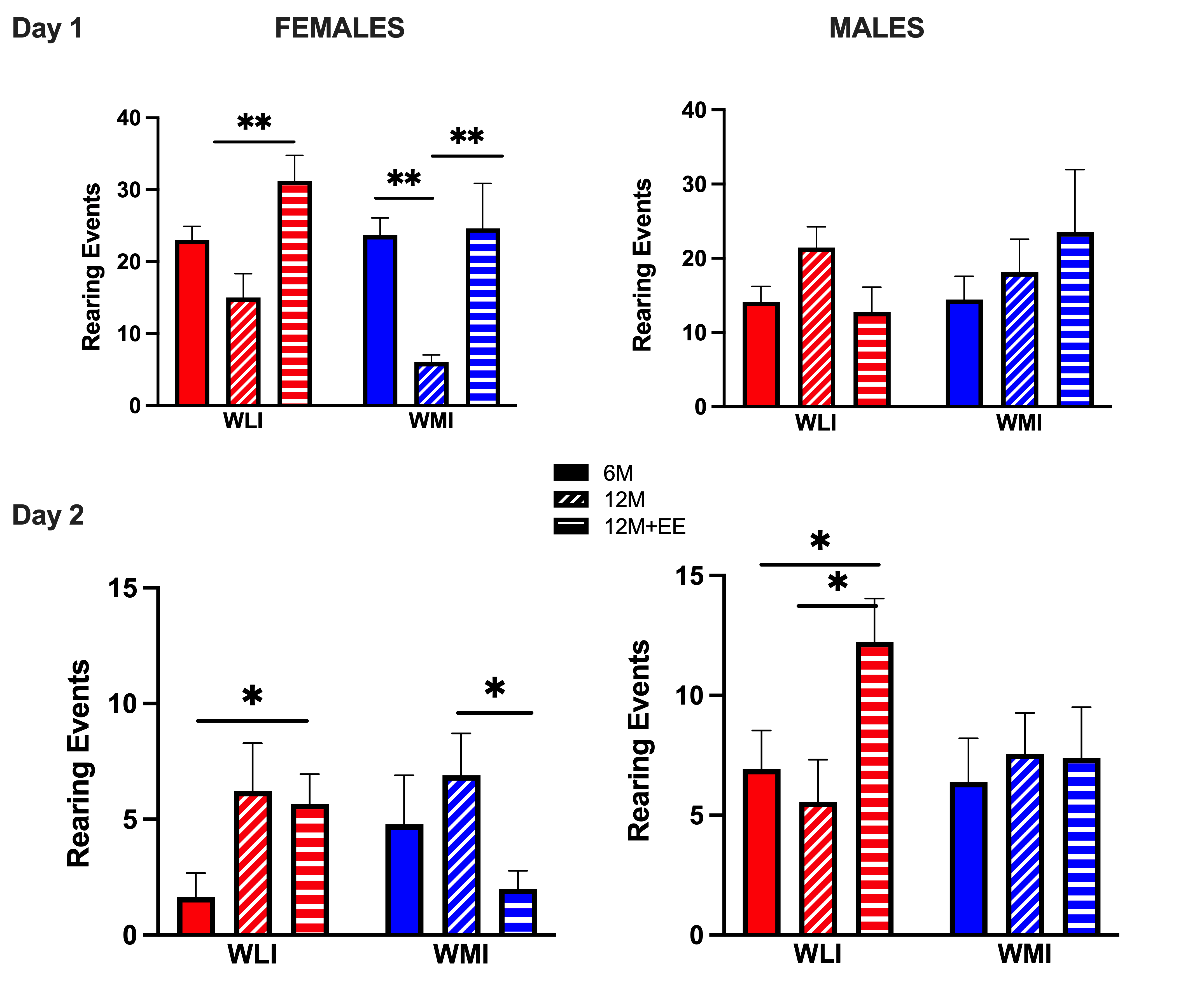

### Supplemental Figure 3

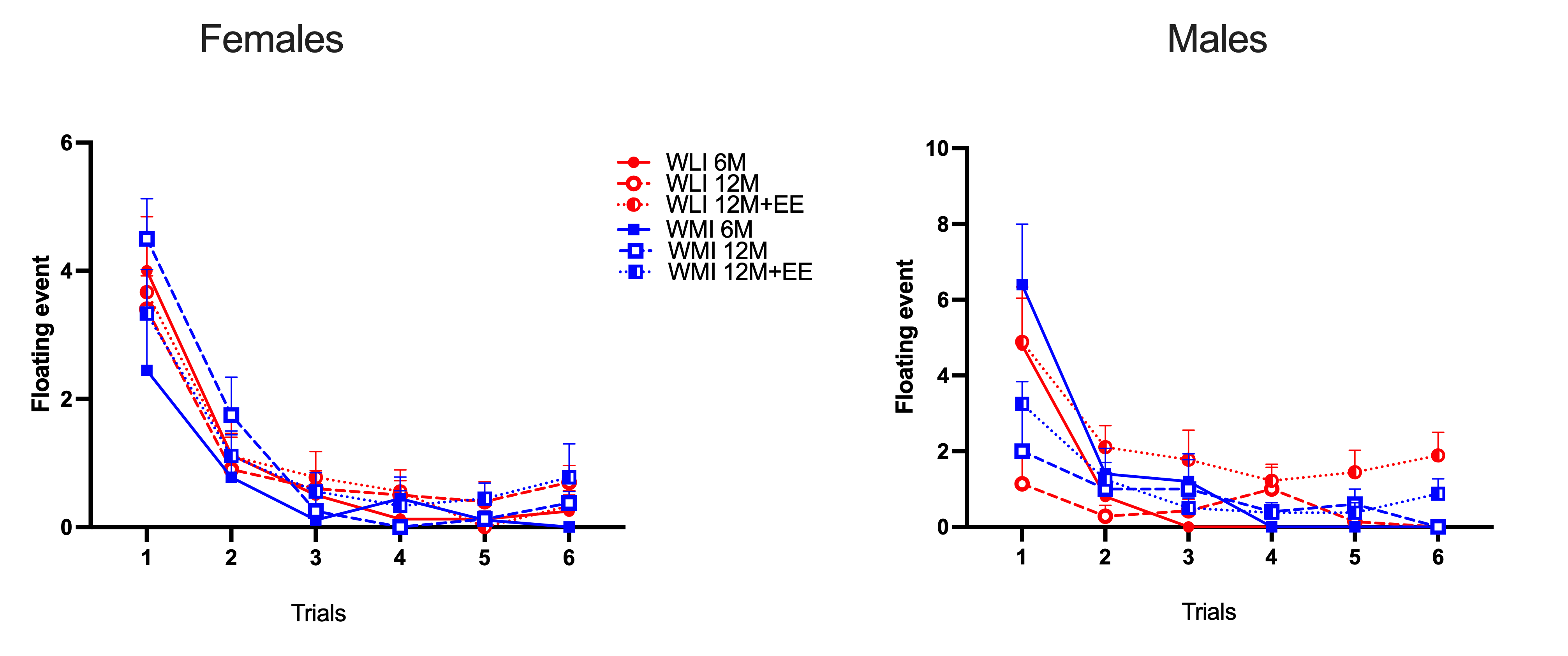

### Supplemental Figure 4

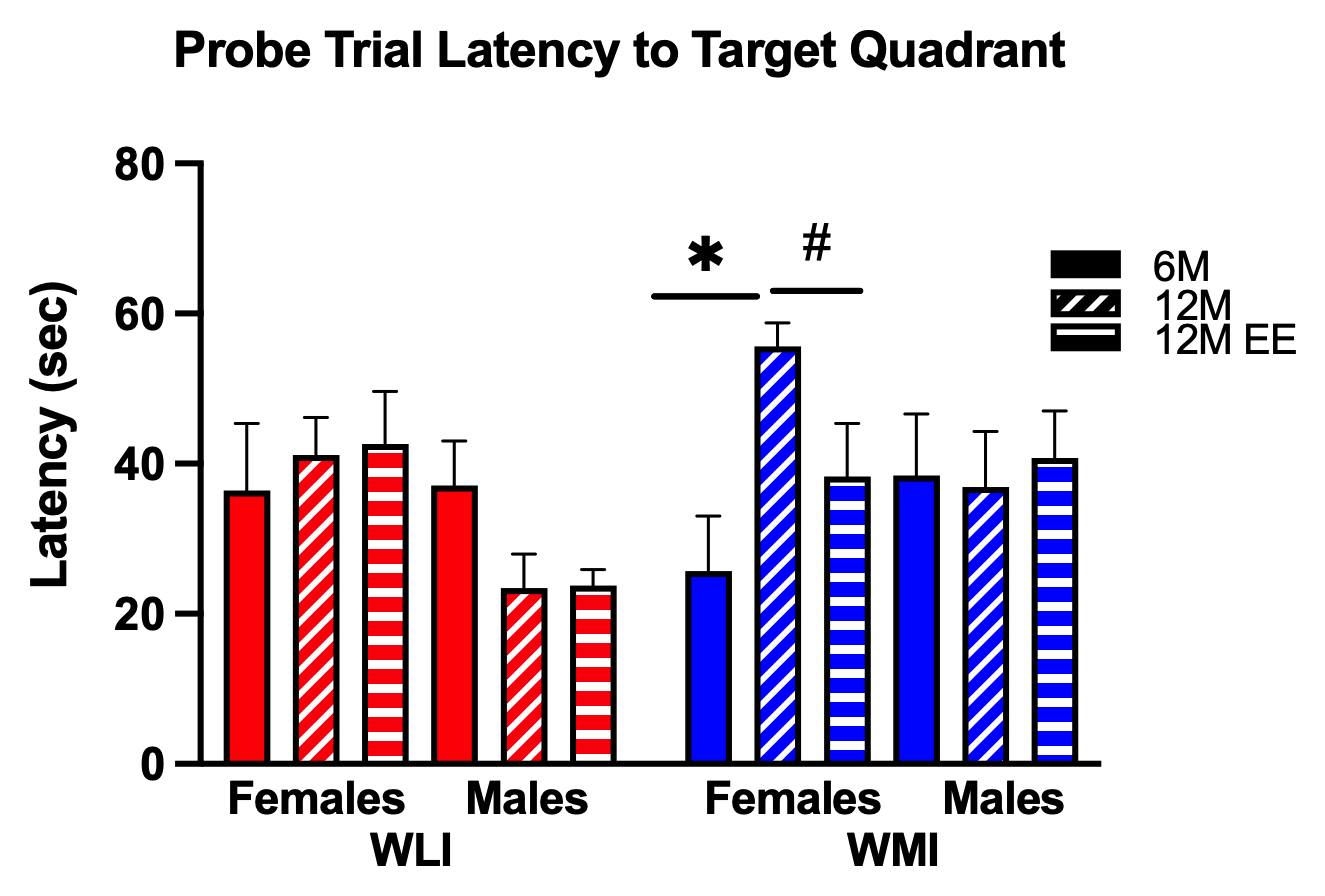
