## Supplemental Table 1 for "Early onset memory deficit of WMI rats compared to their nearly isogenic WLIs is reversed by enriched environment in females"

| **Supplemental Table 1. Primer sequences for quantitative RT-PCR** | | |
| --- | --- | --- |
| **Gene** |  | **Sequence 5’ - 3’** |
| *Mt-co1* | *F* | ATT GGA GGC TTC GGG AAC TG |
|  | *R* | AGA TAG AAG ACA CCC CGG CT |
| *Mt-co2* | *F* | TGG CTT ACA AGA CGC CAC AT |
|  | *R* | TGG GCG TCT ATT GTG CTT GT |
| *Mt-co3* | *F* | AAG GCC ACC ACA CCC CTA TT |
|  | *R* | TAA TTC CTG TTG GGG GTC AGC |
| *Mt-nd3* | *F* | GAC CAA CAA GTT CTG CAC GC |
|  | *R* | AGG GGG AGT AGT AAG GCG AT |
| *Blhle41* | *F* | CGA GAC GAC ACC AAG GA AC |
|  | *R* | AGT GGA ACG CAT CCA AGT CG |
| *Coro2b* | *F* | CCC CCT CCC CTT TTT CCA |
|  | *R* | AGC AGA TCC CAG CCA AAG AC |
| *Slc35a4* | *F* | CTG GGT CAA GTA TGA TGG AG |
|  | *R* | CAA GAG CGA GCC ATC CTG G |
| *Nudt13* | *F* | CAT CGA TAC CTG GTG CCC CG |
|  | *R* | GAA GGAG CTT GAG CCG TGA ACA |
| *Klhl6* | *F* | AGA CCC TGC GAG AGG AAA ATG |
|  | *R* | TAG TTC AGG AGC GTG TGC AT |
| *Smoc2* | *F* | GGT GCC GGA AAA GCA GAT GA |
|  | *R* | CGG CTC TTA ACG TGC TCG GT |
| *Kcnj2* | *F* | TCT CAC TTG CTT CGG CTC AC |
|  | *R* | CCA GAG AAC TTG TCC TGT TGC |
| *Zc3h13* | *F* | AGG GTG CTT AGA GAG GAG TAG T |
|  | *R* | CCA GTT ACG GCA CTG TGT CT |
| *Gapdh* | *F* | CAA CTC CCT CAA GAT TGT CAG CAA |
|  | *R* | GGC ATG GAC TGT GGT CAT GA |
