## Supplemental Table 2 for "Early onset memory deficit of WMI rats compared to their nearly isogenic WLIs is reversed by enriched environment in females"

| **Supplemental Table 2. Strain differences in hippocampal gene expression at 6 months of age (WLI vs. WMI; p<0.01)** | | | |
| --- | --- | --- | --- |
| **Gene name** | **Gene description** | **Fold Change** | **P value** |
| ***Tnfrsf25*** | TNF receptor superfamily member 25 | 0.20 | 0.0052 |
| ***Myh6*** | myosin heavy chain 6 | 0.23 | 0.0052 |
| ***Kcp*** | kielin cysteine rich BMP regulator | 0.26 | 0.0095 |
| ***Wfikkn1*** | WAP, follistatin/kazal, immunoglobulin, kunitz and netrin domain containing 1 | 0.29 | 0.0043 |
| ***Spon2*** | spondin 2 | 0.29 | 0.0074 |
| ***Akr1c19*** | aldo-keto reductase family 1, member C19 | 0.33 | 0.0016 |
| ***Ccn5*** | cellular communication network factor 5 | 0.36 | 0.0001 |
| ***Xkr6*** | XK related 6 | 0.38 | 0.0090 |
| ***Vwf*** | von Willebrand factor | 0.42 | 0.0001 |
| ***Zmynd10*** | zinc finger, MYND-type containing 1NA | 0.42 | 0.0067 |
| ***Acer2*** | alkaline ceramidase 2 | 0.42 | 0.0010 |
| ***Ttll3*** | tubulin tyrosine ligase like 3 [ | 0.43 | 0.0066 |
| ***Hdac7*** | histone deacetylase 7 | 0.46 | 0.0038 |
| ***Adamts6*** | ADAM metallopeptidase with thrombospondin type 1 motif, 6 | 0.46 | 0.0056 |
| ***Kank3*** | KN motif and ankyrin repeat domains 3 | 0.47 | 0.0010 |
| ***Xkr8*** | XK related 8 | 0.50 | 0.0088 |
| ***Pxn*** | paxillin | 0.51 | 0.0007 |
| ***Notch4*** | notch receptor 4 | 0.53 | 0.0012 |
| ***Esam*** | endothelial cell adhesion molecule | 0.55 | 0.0080 |
| ***Doc2g*** | double C2-like domains, gamma | 0.57 | 0.0008 |
| ***AABR07044388.2*** | tenascin XB | 0.57 | 0.0069 |
| ***Stk38*** | serine/threonine kinase 38 | 0.57 | 0.0099 |
| ***Nphp4*** | nephrocystin 4 | 0.59 | 0.0096 |
| ***Cdc45*** | cell division cycle 45 | 0.59 | 0.0047 |
| ***Dennd2b*** | DENN domain containing 2B | 0.60 | 0.0053 |
| ***Fam76b*** | family with sequence similarity 76, member B | 0.60 | 0.0026 |
| ***Hps5*** | HPS5, biogenesis of lysosomal organelles complex 2 subunit 2 | 0.61 | 0.0012 |
| ***Shc4*** | SHC adaptor protein 4 | 0.62 | 0.0016 |
| ***Tsc2*** | TSC complex subunit 2 | 0.63 | 0.0011 |
| ***Parp9*** | poly (ADP-ribose) polymerase family, member 9 | 0.63 | 0.0060 |
| ***Krba1*** | KRAB-A domain containing 1 | 0.64 | 0.0094 |
| ***Mfn1*** | mitofusin 1 | 0.64 | 0.0080 |
| ***NEWGENE_138612*** | MRG domain binding protein | 0.66 | 0.0015 |
| ***Capn15*** | calpain 15 | 0.67 | 0.0011 |
| ***Tbkbp1*** | TBK1 binding protein 1 | 0.67 | 0.0065 |
| ***Slc38a3*** | solute carrier family 38, member 3 | 0.67 | 0.0025 |
| ***Map3k3*** | mitogen activated protein kinase kinase kinase 3 | 0.67 | 0.0005 |
| ***Bicdl1*** | BICD family like cargo adaptor 1 | 0.68 | 0.0015 |
| ***Slc25a42*** | solute carrier family 25, member 42 | 0.69 | 0.0019 |
| ***Scrib*** | scribble planar cell polarity protein | 0.69 | 0.0026 |
| ***Tpcn1*** | two pore segment channel 1 | 0.69 | 0.0016 |
| ***Mcf2l*** | MCF.2 cell line derived transforming sequence-like | 0.70 | 0.0026 |
| ***Dnaja2*** | DnaJ heat shock protein family (Hsp4NA) member A2 | 1.31 | 0.0099 |
| ***Vsnl1*** | visinin-like 1 | 1.32 | 0.0061 |
| ***Mfap1a*** | microfibrillar-associated protein 1A | 1.32 | 0.0046 |
| ***Txnl1*** | thioredoxin-like 1 | 1.33 | 0.0001 |
| ***Ndfip1*** | Nedd4 family interacting protein 1 | 1.34 | 0.0060 |
| ***Mrps2*** | mitochondrial ribosomal protein S2 | 1.35 | 0.0005 |
| ***Snw1*** | SNW domain containing 1 | 1.35 | 0.0065 |
| ***Rcn1*** | reticulocalbin 1 | 1.36 | 0.0065 |
| ***Cadm3*** | cell adhesion molecule 3 | 1.36 | 0.0061 |
| ***Psmc6*** | proteasome 26S subunit, ATPase 6 | 1.36 | 0.0061 |
| ***Lancl2*** | LanC like 2 | 1.38 | 0.0038 |
| ***Cacybp*** | calcyclin binding protein | 1.39 | 0.0020 |
| ***Psmg2*** | proteasome assembly chaperone 2 | 1.40 | 0.0022 |
| ***Snap25*** | synaptosome associated protein 25 | 1.42 | 0.0042 |
| ***Pnma8c*** | PNMA family member 8C | 1.43 | 0.0039 |
| ***Tmx4*** | thioredoxin-related transmembrane protein 4 | 1.44 | 0.0025 |
| ***Ralyl*** | RALY RNA binding protein-like | 1.45 | 0.0015 |
| ***Zbtb33*** | zinc finger and BTB domain containing 33 | 1.50 | 0.0047 |
| ***Kiz*** | kizuna centrosomal protein | 1.54 | 0.0086 |
| ***Kcnc2*** | potassium voltage-gated channel subfamily C member 2 | 1.55 | 0.0010 |
| ***Flrt3*** | fibronectin leucine rich transmembrane protein 3 | 1.62 | 0.0073 |
| ***Qpct*** | glutaminyl-peptide cyclotransferase | 1.98 | 0.0007 |
| ***H4f3*** | H1.3 linker histone, cluster member | 4.58 | 0.0000 |

DEGs are considered when fold change were greater than 1.3 or lesser than 0.7.
