## Supplemental Table 3 for "Early onset memory deficit of WMI rats compared to their nearly isogenic WLIs is reversed by enriched environment in females"

| **Supplemental Table 3. Strain differences in hippocampal gene expression at 12 months of age (WLI vs. WMI; p<0.001)** | | | |
| --- | --- | --- | --- |
| **Gene name** | **Gene description** | **Fold change** | **p value** |
| ***Lcn2*** | lipocalin 2 | 0.30 | 0.0009 |
| ***Smagp*** | small cell adhesion glycoprotein | 0.49 | 0.0006 |
| ***Fmod*** | fibromodulin | 0.56 | 0.0009 |
| ***Pogk*** | pogo transposable element derived with KRAB domain | 0.57 | 0.0009 |
| ***Tmem221*** | transmembrane protein 221 | 0.58 | 0.0001 |
| ***Miip*** | migration and invasion inhibitory protein | 0.60 | 0.0007 |
| ***Smu1*** | SMU1, DNA replication regulator and spliceosomal factor | 1.31 | 0.0009 |
| ***Cct4*** | chaperonin containing TCP1 subunit 4 | 1.31 | 0.0001 |
| ***Ran*** | RAN, member RAS oncogene family | 1.33 | 0.0009 |
| ***Dcaf12l1*** | DDB1 and CUL4 associated factor 12-like 1 | 1.33 | 0.0001 |
| ***Pdhx*** | pyruvate dehydrogenase complex, component X | 1.33 | 0.0009 |
| ***Fezf2*** | Fez family zinc finger 2 | 1.39 | 0.0009 |
| ***Metap1*** | methionyl aminopeptidase 1 | 1.39 | 0.0010 |
| ***Fastkd2*** | FAST kinase domains 2 | 1.42 | 0.0001 |
| ***Bcl2l13*** | BCL2 like 13 | 1.43 | 0.0003 |
| ***Mrap2*** | melanocortin 2 receptor accessory protein 2 | 1.51 | 0.0004 |
| ***Ipcef1*** | interaction protein for cytohesin exchange factors 1 | 1.52 | 0.0008 |
| ***Kcnab3*** | potassium voltage-gated channel subfamily A regulatory beta subunit 3 | 1.69 | 0.0002 |
| ***Gna14*** | G protein subunit alpha 14 | 1.92 | 0.0002 |
| ***Mgat4c*** | MGAT4 family, member C | 1.99 | 0.0011 |
| ***Morc3*** | MORC family CW-type zinc finger 3 | 2.08 | 0.0003 |
| ***Brinp3*** | BMP/retinoic acid inducible neural specific 3 | 2.13 | 0.0007 |
| ***Cntn5*** | contactin 5 | 2.34 | 0.0010 |
| ***Rspo3*** | R-spondin 3 | 2.41 | 0.0003 |
| ***Sgpp2*** | sphingosine-1-phosphate phosphatase 2 | 2.71 | 0.0011 |
| ***AABR07006724.1*** | protein kinase cGMP-dependent 1 | 2.80 | 0.0010 |
| ***Cdh12*** | cadherin 12 | 2.91 | 0.0001 |
| ***Bmp3*** | bone morphogenetic protein 3 | 3.09 | 0.0009 |
| ***H4f3*** | H1.3 linker histone, cluster member | 6.45 | 0.0008 |
